# A novel dopaminergic critic signal triggered by erroneous strategy choices in mice training in operant tasks

**DOI:** 10.1101/2024.05.06.592735

**Authors:** Jumpei Matsumoto, Virginie J. Oberto, Marco N. Pompili, Ralitsa Todorova, Francesco Papaleo, Hisao Nishijo, Laurent Venance, Marie Vandecasteele, Sidney I. Wiener

## Abstract

Classically, midbrain dopaminergic neuron activity is triggered by unexpected rewards, then, upon learning, by reward-predictive conditioned stimuli. When expected rewards are withheld, firing is inhibited. This activity occurs too late to directly affect the neuronal circuitry underlying decision-making, inspiring the development of temporal difference (TD) reinforcement learning models. To test for more timely critical feedback during decision-making and learning, we recorded optogenetically identified dopaminergic, putative GABAergic and other neurons of the ventral tegmental area (VTA) and substantia nigra pars compacta in mice training in visual and olfactory discrimination tasks. The mice often adhered to unrewarded and untrained task strategies (e.g., spatial alternation) rather than making random choices. In order to probe for reward/punishment predictive activity, a delay was imposed between nose-poke choices and signals for reward or punishment. As animals performed below criterion levels, dopaminergic and other neurons’ firing rates signaled correct versus incorrect choices immediately after choices, but prior to the onset of trial outcome signals. Thus, this activity signaled the rewarded rule even as the mice performed other unrewarded strategies. Putative GABAergic neurons fired during nose-poke choices, potentially reducing network activity prior to reward prediction signals. This reward predictive activity could serve as a critic signal expressed immediately after choices are made, priming the network for canonical DA reward/punishment activity, facilitating network functional modifications. This is consistent with a role for dopamine in arbitration between brain modules to choose among diverse strategies during goal-directed behavior. These findings suggest extensions of theoretical formulations interpreting dopaminergic neuronal activity.

**Significance statement:** Models of dopaminergic influence on circuit modifications during learning evoke mechanisms dealing with the delay between neural activity leading up to choices, and when the reinforcing outcome actually occurs. Here, as mice trained in sensory discrimination tasks with a delay between behavioral responses and reinforcement signals, they performed several unrewarded behavioral strategies. Simultaneously, dopaminergic nuclei neurons instead reflected the current task rule, predicting whether the choice was correct or not, providing an immediate “critic” signal prior to canonical trial outcome signals. This provides evidence for brain mechanisms to overcome innate or acquired habits to perform behaviors optimizing positive outcomes.

## Introduction

Behavioral adaptation for survival requires associating predictive cues within their contexts with appropriate behavioral policies to favor positive outcomes at low cost and to avoid aversive ones. Dopaminergic activity is strongly implicated in the modification of neural circuits making such associations. Early in operant training, dopaminergic neurons fire phasically after unexpected rewards (Hollerman & Schultz, 1998). The DA neuron activity after rewards persists on trials immediately following learning acquisition, and then gradually diminishes on subsequent trials. Following learning, spiking activity is triggered by presentations of a conditioned stimulus (CS) that predicts the reward, while inhibition occurs when the expected reward is withheld.

Such activity is interpreted as evidence that dopaminergic neurons signal “reward prediction errors” (RPEs) (Houk et al., 1995; Montague et al., 1996, Schultz et al., 1997). RPE activities are posited to code motivational value (Schultz et al., 1997; Wise, 2005) to correspond to a neural substrate for the “temporal difference (TD)” machine-learning algorithm which would modify the circuitry underlying behaviors leading up to these outcomes. To deal with the problem that this neural activity may occur at long delays prior to the outcome, TD learning estimates the expected reward on the basis of the current “state”, that is, the ensemble of environmental context and cues, as well as the agent’s actions. The expected reward is compared between the current state, and preceding estimates. Any differences, positive or negative, referred to as the “TD error”, can then be employed to improve estimates of the state value. This “critic” dopaminergic signal would then modify activity of circuits including striatal “actors” to enact adaptive action selection (e.g., Khamassi, et al., 2005).

Interestingly, in basolateral amygdala and orbitofrontal cortex recordings, Schoenbaum, et al. (1998, 1999) observed neurons that discharged in the interval between choice and outcome signals and that their activity distinguished correct from incorrect choices. Recently, Cazettes et al. (2023) recorded neurons in M2 cortex (which projects to ventral tegmental area, VTA, and substantia nigra pars compacta, SNc; Watabe-Uchida et al., 2012) in mice performing tasks with multiple possible response policies, and observed representations of unused strategies. Ding et al. (2025) observed reward prediction in population activity of putative DA neurons in VTA after correct choices prior to reward. Lak, et al (2020) recorded VTA dopamine neuron activity using fiber photometry of GCaMP6 signals. They imposed delays between moving optic grating stimuli and signals for the mice to respond by turning a wheel and observed increases in activity for stimuli leading rewards. While that study focused on varying visual cue saliency, the present work examines dopaminergic activity as mice select strategies in tasks depending upon cues of different modalities in the presence and absence of distractors. We hypothesized that when the behavioral response policy can be selected among a variety of behavioral strategies, a subset of VTA neurons could also represent the rule currently being acquired during training, even while the animals are performing other strategies, since this could eventually serve as a timely critic signal prior to reward or punishment.

To test for such activity, we recorded mice as they trained in and acquired odor discrimination and visual discrimination tasks with and without distractors successively within the same experimental chamber. Once criterion performance was achieved, the task was modified, requiring continual adaptation to new contingencies. Trial outcome cues (signaling reward or punishment) were delayed until after the animals made nose-poke choices in order to facilitate detection of outcome prediction signals. The mice were a mutant strain, permitting optogenetic identification of dopaminergic neurons with light stimulation (Eshel et al., 2015). The animals often executed a variety of unrewarded strategies prior to ultimately reaching criterion performance. As anticipated, we observed a novel pattern of phasic dopaminergic neuron activity in the 0.5-1.0 s interval after nose-poke choices but prior to trial outcome signals. Thus, this activity signaled future reward or punishment immediately following decisions. Most of these same neurons also had RPE activity: excitation following to reward delivery signals, and inhibition after punishment signals (Schultz, et al., 1997). A theoretical framework is presented to illustrate a possible role for these dopaminergic activities in deciding among multiple strategies and reinforcement learning.

## Materials and Methods

### Animals

All experiments were performed in accordance with EU guidelines (Directive 86/609/EEC). Subjects were four transgenic adult (2-9 months) male mice expressing a channelrhodopsin2-yellow fluorescent protein fusion protein (ChR2 (H134R)-eYFP) on dopamine neurons. These mice were obtained by mating mice expressing the CRE recombinase under the control of the dopamine transporter (DAT-IRES-CRE mice, Stock 006660, Jackson Laboratory, ME, USA) with mice bearing a CRE-dependent ChR2(H134R)-eYFP gene (Ai32 mice, Stock 012569, Jackson Laboratory). Animals were housed in temperature-controlled rooms with standard 12-hour light/dark cycles and food and water were available ad libitum. Each workday, animals were handled to habituate to human contact, and weighed. During pre-training and experimental periods, food intake, including pellets provided in the experimental apparatus, was restricted to a maximum of 3 g/day. Water access remained ad lib. Supplemental food was provided if weight fell below 85% of the normal weight.

### Surgery

Anesthesia was induced with a mixture of ketamine (66 mg/kg) and xylazine (13 mg/kg) and sustained with isoflurane (0.5 – 1.0%). Mice were placed in a stereotaxic device (David Kopf Instruments) and maintained at 37° C. The scalp was exposed and cleaned with hydrogen peroxide solution. Miniature jeweler’s screws were screwed into trephines, and attached with dental cement. A custom electrode assembly (developed by JM; see Oberto, Matsumoto, et al., 2023) was implanted into the left VTA and SNc. Briefly, the assembly consisted of microdrives holding four recording probes for recording neural activity and two optic fibers for optogenetic identification with light stimulation. Each recording probe was twisted bundles of 8 formvar coated 12-micron diameter nichrome wires (“octrodes”) were inserted in polyimide tubes. The wires were gold-plated to an impedance of about 350 kohm. The stainless-steel screws implanted on the left and right cerebellum as ground and reference electrodes, respectively.

### The behavioral task

#### Experimental apparatus

The ‘Operon’ system developed by Scheggia et al (2014) was adapted as an experimental chamber permitting the mice to perform olfactory and visual discrimination tasks (Figure 1). Many components were purchased from Med Associates (Fairfax, VT, USA). Each of the short sides of the rectangular (160L x 136W x 160H mm) plexiglass chamber has two arrays of LEDs and two ports used for both odor sampling and nose pokes. An olfactometer presented d- and l-stereoisomers of limonene at the respective odor ports in a pseudo-random sequence. (In a few sessions, two other odors, pine and citronella, were used.) All six white LEDs were lit above one nose-poke port and remained unlit above the other, again in pseudo-random sequence. Between the ports is a central feeder port with an overhead lamp (“house light”) which was lit for 7 s during the delay after error trials. This lamp emits only 55 lux, which is not intrinsically aversive. As a reward, a pellet (5TUL TestDiet, Richmond, IN, USA, 14 mg) was delivered by a dispenser which made an audible sound (ENV-203-14P, Med Associates, Fairfax, VT, USA). A plexiglass barrier divided the two sides of the chamber and was slid up from below via an automated motor assembly. Behavioral procedures were adapted from those of Scheggia and Papaleo (2016) but without textural cues, and further details can be found there and in Scheggia et al. (2014). Lights in the experimental room were turned off for recordings.

**Figure 1.**
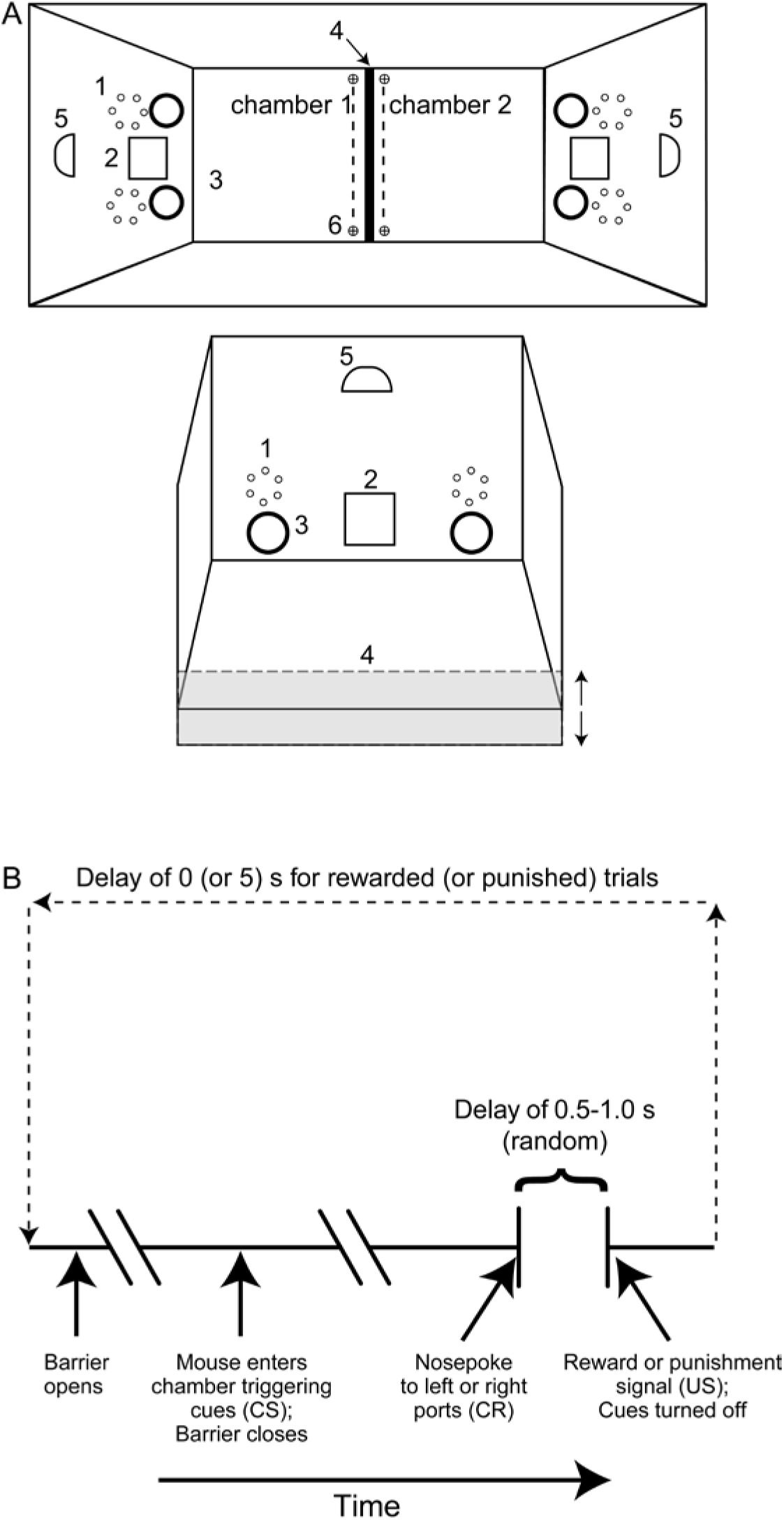
The behavioral apparatus and task (after Scheggia, et al., 2014). **A)** Top) Overhead view. 1- Hexagonal LED array; 2- Reward dispenser; 3- Nose-poke port; 4-Moveable barrier; 5-House light; 6- Photobeam. Bottom) Frontal view of one chamber (photobeam not shown here). **B)** The automated task sequence.

#### Pre-training

Mice were provided water ad libitum, but mildly food deprived to maintain them above 85% their baseline weight, as controlled with daily weighing. The behavioral apparatus was turned on, and the olfactometer outputs were confirmed to present odors discriminable by the experimenter. The room lights were dimmed and the behavioral task was started. Then, with the barrier up, the mouse was placed into one of the compartments. To signal the beginning of each trial, the barrier was lowered. Once the mouse crossed the central photodetectors on its way to the other the chamber, one of the two LED arrays was lit and the two odors were released from the respective odor ports in that chamber. This photodetector crossing is called the “Cue” event. Once the mouse cleared the central photodetectors, the barrier was raised. Nose-pokes to the port on the side of the (currently rewarded) odor or visual cue blocked a photodetector within the port, and, at a variable latency ranging from 0.5 to 1.0 s, this triggered trial outcome signals. Choices adherent to the current rule triggered the audible release of a food pellet (“Reward”). Erroneous choices triggered illumination of an overhead light during a timeout period of 5 or 8 s while the barrier remained up (“Punishment”). Animals were first pre-trained to shuttle between two compartments for reward. In the first stage of training, a nose poke into either port triggered a pellet delivery. Then animals were pre-trained for simple visual discrimination task (LED’s on vs LED’s off). Once they could regularly perform sequential trials in the maze, mice were returned to their cages and provided with food ad lib for at least several days prior to implantation surgery.

#### Behavioral tasks during recordings

After recovery from optrode assembly implantation, the mice were first recorded as they re-acquired the “simple” visual discrimination task. Once mice attained criterion performance, the system automatically triggered a change in task contingencies. (These criteria were selected to prevent overtraining and lengthy persistence in the learned strategy. The justification for selection of criterion values for performance are discussed below.) Typically, after acquisition of the simple visual discrimination task, the olfactory cues were also emitted from the respective nose-poke ports when the visual cue was lit. These olfactory cues served as distractors, and thus this is termed the “complex” visual discrimination task. Once criterion performance was again reached in the complex visual discrimination, the reward contingency was shifted to the olfactory discrimination task, either the simple version with no visual distractor cues, or the complex version with the visual distractor cues. When the mice reached the criterion performance level, the reward contingency was again shifted. In general, each session started with a simple (visual or odor) discrimination task, then proceeded to the complex version prior to shifting to discriminations with the other modality. Sometimes when mice seemed to be performing poorly, the complex task was reverted to a simple one. When the mouse stopped responding for more than 120 s, the behavioral session was ended. The weight of the rewards earned was calculated, and the remaining required amount of chow for the daily feeding was provided.

Each series of trials with visual or olfactory discrimination in simple or complex versions is referred to below as a “task epoch”, and these periods are analyzed separately since a given neuron could respond differently in different task epochs (as we previously found in the VTA/SNc-afferent nucleus accumbens; Shibata, et al., 2001 and will be shown below). The extra-dimensional attentional set-shifting nature of this task (Scheggia, et al., 2014) was reported to require intact cingulate cortical function (Dias, et al., 1996; Birrell & Brown, 2000; Bissonette, et al., 2008), which, in turn, depends upon dopaminergic input (see e.g., Vander Weele, et al., 2019; Di Domenico & Mapelli, 2023).

The position of the rewarded cue was pseudo-randomly varied on successive trials so that the same odor or visual cue were not presented more than twice at the left or the right nose-poke port. While the rewarded side was varied pseudo-randomly, in exceptional cases when an animal persisted at visiting one side, the other side was rewarded more. Furthermore, the same average trajectory orientation across the chamber (diagonally crossing between the two chambers or running along a side wall) was not rewarded on more than 2 successive trials.

### Justification of the choice of strategies analyzed and for selection of criterion values for performance

In previous work, we searched for most likely alternative strategies on the basis of a Bayesian descriptive model of behavioral data in a comparable maze (Peyrache et al., 2018) which tested possible 256 rule combinations (Annex of Khamassi, et al., 2024). The latter study concluded that the choice of the rules Left port preference (or Right), Lit port preference (or Unlit) and Alternate left and right choices (or its alternative, Persist at left or right choices) were optimal for describing the rats’ behavior. There was no odor discrimination in that study, and hence odor strategies were added here. Since Persist and Left (or Right) have overlapping definitions, the Persist strategy was not considered further.

The task rule (i.e., reward contingencies) was changed between simple and complex and/or between modalities when the mice had received eight rewarded choices out of ten consecutive trials, which typically included 6 consecutively rewarded trials; see Figure 2). This criterion was intended to avoid overtraining and sustained persistence at a given strategy. Furthermore, the probabilistic analyses of behavioral data from a similar task in the Annex of Khamassi et al (2024) support the choice of the criterion of six successive compliant trials as a criterion for describing adherence to a rule and excluding fortuitous inclusion of series of rewarded stochastic choices. Note that these criteria are consistent with those typically used in this type of study (e.g., Kaefer, et al., 2020; Lapiz-Bluhm, et al., 2009, Tait, et al., 2017).

**Figure 2.**
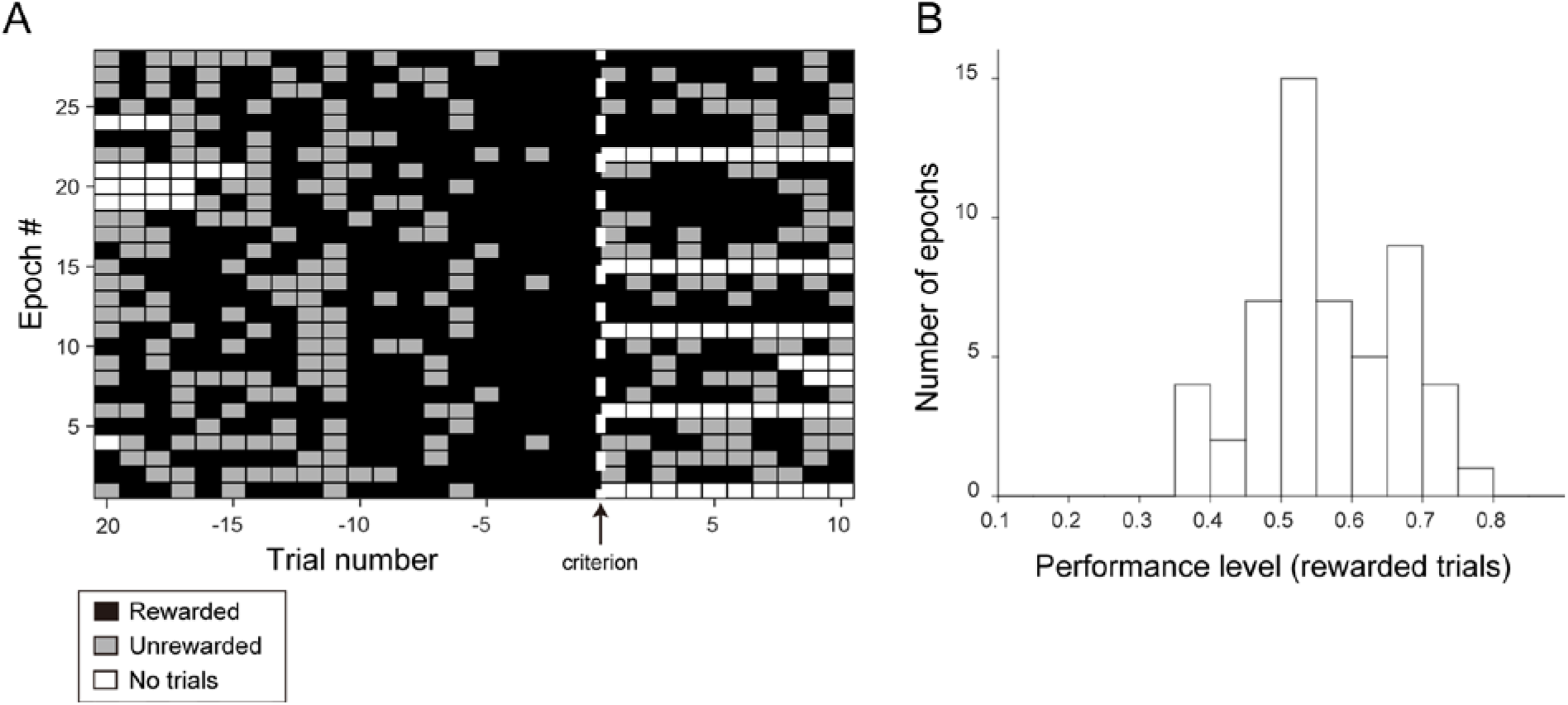
Task performance. A) Profile of rule acquisition in trials leading up to reaching the criterion for changing task rules (80% rewarded) in representative epochs where DA reward predictive neurons (described below) were recorded. “No trials” correspond to before the beginning or after the end of the session. B) Distribution of mean performance levels (proportion of trials that were rewarded) in representative epochs with reward-predictive DA neurons. N = 54 epochs. If more than one responsive cell was recorded in a given epoch, the epoch was only counted once.

Slightly different criteria were applied for performance criteria in analyses of incidences of unrewarded strategies during training. Again, these were selected on the basis of Bayesian analyses of simulations of random choices in the behavioral data set of Khamassi et al. (2024). For the analyses of incidences of series (or “runs”) of trials with unrewarded strategies, two approaches were adopted. The “strict adherence” approach did not permit any non-adherent trials in the run. For the more “permissive adherence” approach, intermittent single non-adherent trials were allowed, provided that they were before or after at least six adherent trials. Thus, sequences of at least 6 (or 8) trials satisfying these criteria were retained. In cases where successive series contained overlapping trials adhering to two different strategies, the overlapping trials were excluded.

### Recording procedure

Recordings of neural signals were made with an Ampliplex system (sampling rate: 20 kHz; cutoff frequencies of the analog low-pass and high-pass filters were 0.3 and 10 kHz, respectively). After surgical implantation, electrodes were advanced as often as twice daily until reaching dopaminergic neurons responsive to optical stimulation. Electrodes were further advanced when discriminable units were no longer present.

On each recording day, the headstage was plugged in and the mouse was placed in the behavioral apparatus. Then the neural signals were recorded as the mouse performed tasks. The position of a LED attached to the headstage was captured with a video camera (30 frames/sec) installed above the system. After the mouse stopped performing the task, the headstage plug was changed, optic fibers were connected, and the animal was placed in a large plastic beaker. Using a 475 nm-laser light source (DPSSL BL473T3-100FL, Shanghai Laser and Optics Century, Shanghai, PRC), trains of 10 light pulses (pulse duration: 10 msec; light power: 10 mW max; frequency: 5 Hz) were given for 20 times with 10 sec inter-train intervals under control of an Arduino-based system designed by JM.

### Data analyses

#### Spike sorting and neuron type classification

For single unit discrimination from the extracellular signals recorded from the octrodes, offline spike sorting was carried out with KiloSort (Pachitariu et al., 2016) followed by manual curation using Klusters (L. Hazan, http://neurosuite.sourceforge.net).

To identify neurons as dopaminergic (DA), we used the Stimulus-Associated spike Latency Test (SALT; Kvitsiani et al., 2013; Eshel et al., 2015). This test determines whether light pulses significantly changed a neuron’s spike timing by comparing the distribution of first spike latencies relative to the light pulse, assessed in a 10-msec window after light-stimulation, to 10 msec periods in the baseline period (-150 to 0 msec from the onset of light-stimulation; see Kvitsiani et al., 2013 for details). A significance level of p<0.01 was selected for this. All neurons referred to below as DA passed this test.

All neurons recorded from an octrode with at least one SALT+ response were considered to be within the dopaminergic VTA or SNc. If the octrode was not advanced, neurons on the day before and the day after were also counted as in VTA or SNc, even if no SALT+ responses were recorded on those days. Similarly, if the octrode had not been advanced, and SALT+ responses had been recorded on non-consecutive days, intervening days with no SALT+ responses were still considered as in a midbrain dopaminergic nucleus. All neurons qualifying as in dopaminergic nuclei were categorized with clustering according to spike width and mean firing rate (see Figure 3). The criterion for putative GABAergic (pGABA) neurons was firing rate > 15 Hz and spike width < 1.5 ms (cf., Ungless and Grace, 2012). No pGABA neurons were SALT+. All other SALT- neurons in dopaminergic nuclei are referred to as “Unidentified”.

**Figure 3.**
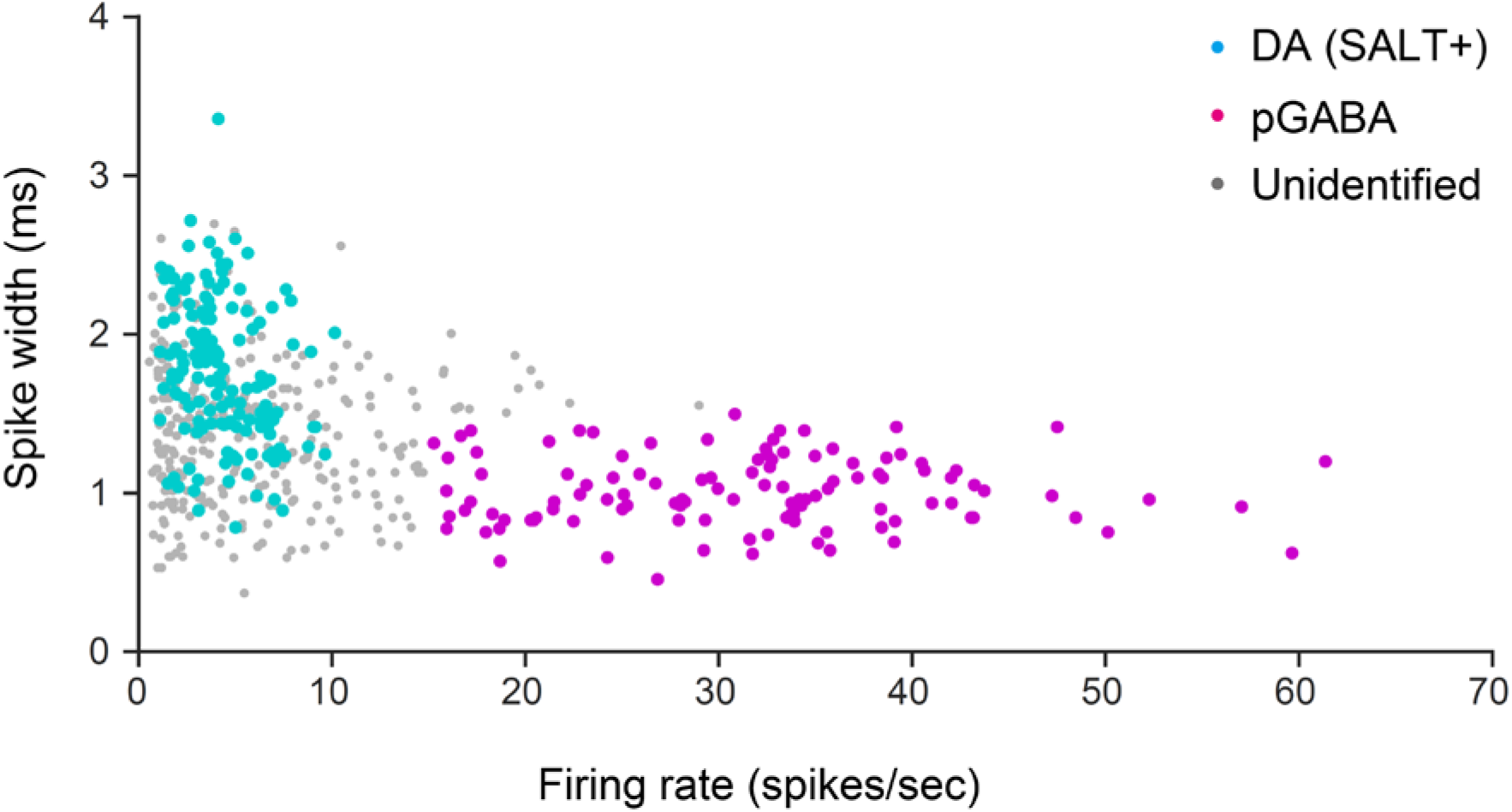
Characterization of neuron identities.

#### Behavioral correlates of neural activity

Neurons with mean firing rates <1 spike/s in a given epoch were excluded from analyses. The Arduino-based task control system (developed by JM) permitted programming the task contingencies with appropriate reward or punishment delivery, outputting time stamps for cue onsets, nose-poke responses, and outcome (reward or punishment) cue onsets. For each neuron in each epoch, firing rates were calculated for baseline periods ([cue onset-0.5 s, cue onset]), [cue, nose-poke] intervals, [nose-poke, outcome] intervals, and post-outcome intervals ([outcome signal, outcome signal+0.5 s]). The RP responsive neurons were defined as those showing significant difference(s) between rewarded and non-rewarded trials in either or both of [cue, nose-poke] and [nose-poke, outcome] intervals (unpaired t-test, p < 0.05) over the course of a task epoch. Similarly, the activity changes after reward (or punishment) signals were tested by comparing firing rates in post-outcome intervals of rewarded (or punished) trials with the baseline period (unpaired t-test, p < 0.05).

### Experimental design and statistical analyses

Neurons were recorded in four transgenic adult (2-9 months) male mice expressing a channelrhodopsin2- yellow fluorescent protein fusion protein (ChR2 (H134R)-eYFP) on dopaminergic neurons. In order to respect “Reduction” of the 3R’s, experiments were stopped when sample sizes were decided to be sufficient, i.e., when significant effects could be determined reliably and replicated.

Basic statistical methods such as paired and unpaired t-test, and Pearson’s correlation analysis were performed using MATLAB. Chi-squared tests were performed with the online calculator at www.socscistatistics.com/tests/chisquare2/calculator/. Significance thresholds were set to p<0.05 unless stated otherwise. The values for n, p, and the specific statistical test performed for each analysis are included in the corresponding figure legend, table, or the main text.

### Histology

Once stable recordings were no longer possible, marking lesions were made with 10 s of 30 µA cathodal current. After waiting for at least 90 min, mice were then killed with a lethal intraperitoneal injection of sodium pentobarbital, and perfused intra-ventricularly with phosphate buffered saline solution (PBS), followed by 10% phosphate buffered formalin. The brain was removed, post-fixed overnight, and placed in phosphate buffered 30% sucrose solution for 2-3 days. Frozen sections were cut at 80 µm, and permeabilized in 0.2% Triton in PBS for 1 h at room temperature. Antibody staining was performed to confirm localization of electrodes in dopaminergic nuclei. Sections were then treated with 3% bovine serum albumen (BSA) and 0.2% Triton in PBS for 1 h with gentle agitation at room temperature to block non-specific binding. Sections were then rinsed for 5 min in PBS at room temperature with gentle agitation. Then the sections were left overnight with gentle agitation at 4° C in a solution of the first antibody, mouse monoclonal anti-TH MAB318 (1:500), 0.067% Triton, and 1% BSA in PBS. After three rinses for five minutes in PBS, the sections were treated with a second antibody (1/200, anti-mouse, green), Nissl-red (1:250), and 0.067% Triton in PBS for two hours at room temperature. Then sections were rinsed 3 times for five minutes in PBS, and mounted with Fluoromount®. Sections were examined with fluorescence microscopy to verify electrode tips and trajectories in the immunofluorescent stained zones.

### Behavior

Neurons were recorded in four mice performed as they performed 14167 trials in 386 task epochs in 167 recording sessions. An epoch is defined as a series of trials with one of the four tasks: visual or olfactory discriminations with a distractor cue (“complex” version) or without (“simple”). Once the mouse reached a criterion of at least eight out of ten successive trials rewarded, the rule was usually changed in order to avoid overtraining and persistence (see Materials and Methods). The mice learned the tasks (examples in Figure 2A). Mice often reached criterion within 40 trials or less (Figure S1). On the other hand, despite extensive experience (thousands of training trials in previous sessions), there remained epochs with more than 50 trials when criterion was not reached (Figure S1). Because of these delays and the task rule changes upon reaching criterion, the mice received rewards in only 56 ± 1% of the trials (mean ± SEM; Figure 2B). Since each session continued until the animal stopped performing, criterion performance was rarely reached in the final epoch.

Since the primary focus here is on DA neuron activity during goal-directed reinforcement learning and associated decision-making, it was important to verify whether behavioral performance actually improved during recording sessions. Indeed, the mice ultimately achieved rule-shift criterion performance in the task in 28 of the 54 task epochs where one or more optogenetically identified DA neurons with reward-predictive activity (characterized below) were recorded (Figure 2A). In 15 other epochs of these 54, the mice also achieved this criterion performance, but in another epoch in the session.

### Mice followed unrewarded strategies during rule acquisition

Interestingly, as the animals performed poorly at adhering to the currently rewarded rule and many choices were not random. During all recording sessions, animals performed at strategies other than the current rule on over 25% of the trials (Figure 4).

**Figure 4.**
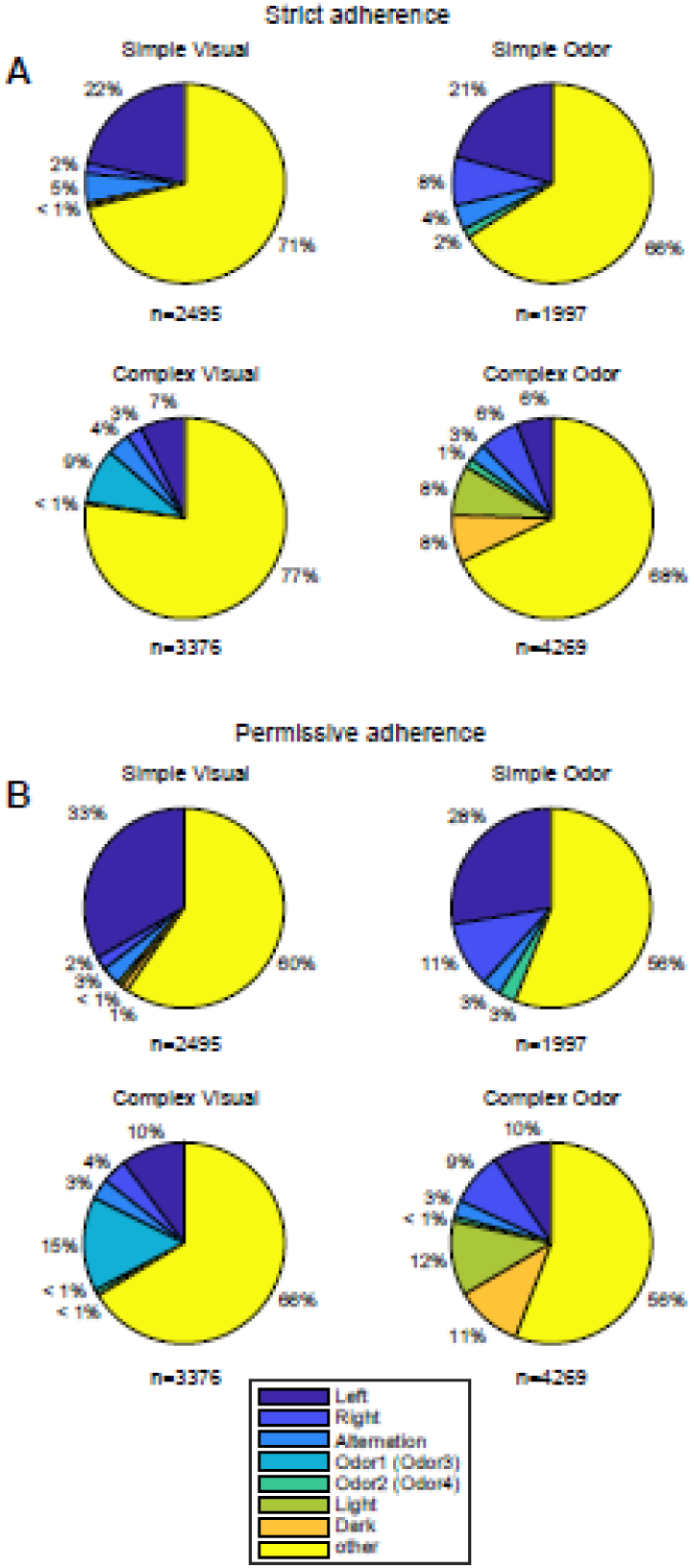
Incidence of adherence unrewarded strategies in the four tasks for all recording sessions. (These criteria are justified in the Materials and Methods section). The “other” category includes undetermined strategies as well as trials adhering to the currently rewarded rule prior to reaching the performance criterion. Only epochs with 15 or more non-criterion performance trials were tallied.

### Neuronal recordings Reward prediction activity

In 167 recording sessions in four mice, of a total of 967 neurons recorded, 272 were optogenetically identified as dopaminergic (DA) neurons, and 206 were putative GABAergic neurons (pGABA), while 489 others did not qualify as DA or pGABA. The latter are referred to as “Unidentified” neurons (Figure 3; see Materials and Methods).

While mice were performing at sub-criterion (Figure 2B) levels in the sensory discrimination tasks, neurons fired selectively according to whether the trials would be rewarded or not, but prior to cues signaling that a reward or punishment was imminent. These are referred to as “reward prediction” (RP) activity. Anatomical reconstruction with fluorescence microscopy revealed that RP responsive neurons were recorded within the VTA and SNc at mediolateral positions ranging from 317 to 1394 µm (see Supp. Fig. 3D of Oberto, Matsumoto, et al., 2023). Analyses were first performed for the [cue, outcome signal] interval. This selectivity occurred in 82 (13%) of the DA neurons recorded in 637 task epochs, 109 Unidentified neurons recorded in 974 epochs (11%) and 31 pGABA neurons in 411 epochs (8%). Among the RP responding neurons, significant increases in firing rate predicting rewards (referred to as “REW>PUN”) had a different incidence than increased firing prior to punishments (PUN>REW) among the three cell groups: 85% were REW>PUN for DA, 55% for Unidentified, and 68% for pGABA (χ^2^(4) = 28.1, p = 1.2E-5; Table 1). The REW>PUN RP activity occurred more frequently in DA neurons than the other two cell types (χ^2^(2) = 16.7, p = 2.4E-4; Table 1). To estimate the magnitudes of significant RP activity, an “RP index” was calculated from firing rates (FR) in the [cue, choice] and [choice, outcome signal] intervals as (rewarded FR - punished FR)/(rewarded FR + punished FR). These are shown in Figure S2.

In these mice, RP activity was detected in the odor discrimination task at the earliest after 84 trials (most took longer) and after the mouse reached criterion performance at least one time. (This could not be calculated for the visual discrimination task since mice were pre-trained in this task prior to optrode implantation. It is possible that there was RP activity earlier, but activity from such neurons were not sampled.) Scheggia et al. (2016) studied mice not equipped with electrode prostheses in similar version of this maze, and they achieved criterion performance (eight rewarded out of ten consecutive trials) in the simple visual and the simple odor discrimination tasks after 60 training trials on average. This is consistent with these strategies being acquired, not innate preferences.

To determine whether RP activity developed gradually over the course of the epoch, the RP index were computed for the first six rewarded and unrewarded trials of the epoch and compared with the last six pairs of trials. For REW>PUN RP activity, the increases in RP indices were significant for DA neurons (p=0.022, Wilcoxon signed-rank test) but not pGABA and Unidentified neurons (p=0.16 and 0.10). For PUN>NR, DA and pGABA RP index decreases were not significant (p=0.098 and 0.38 but they were for Unidentified neurons (p=0.035; Figure S3). Despite these significance trends in the population data, results were diverse for individual epochs (Figure S3).

### DA neurons

The onset of RP activity is an indicator of the latest moment in the trial when the behavioral choice could have been determined in the brain. In the [cue, nose-poke] interval, 10% of the epochs had significant REW>PUN responsive DA neurons, while in the [nose-poke, outcome] interval, 18% were significant (Table S1). For PUN>REW, these values were only 2% and 3%, respectively. REW>PUN RP activity in DA neurons primarily occurred in the [cue, nose-poke] interval when the target cue (i.e., associated with the current reward contingency) was visual (82%) rather than olfactory (Figure 4A, “Difference”; Table S1). In contrast, in the [cue onset, nose-poke] interval, olfactory cues were the target in 81% of the REW>PUN RP epochs (Figure 5A, Table S1). One possible explanation for the different timing of onset of RP activity in the two tasks would be if the visual cues were detected and discriminated earlier than the olfactory cues, and the behavioral response was decided then.

**Figure 5.**
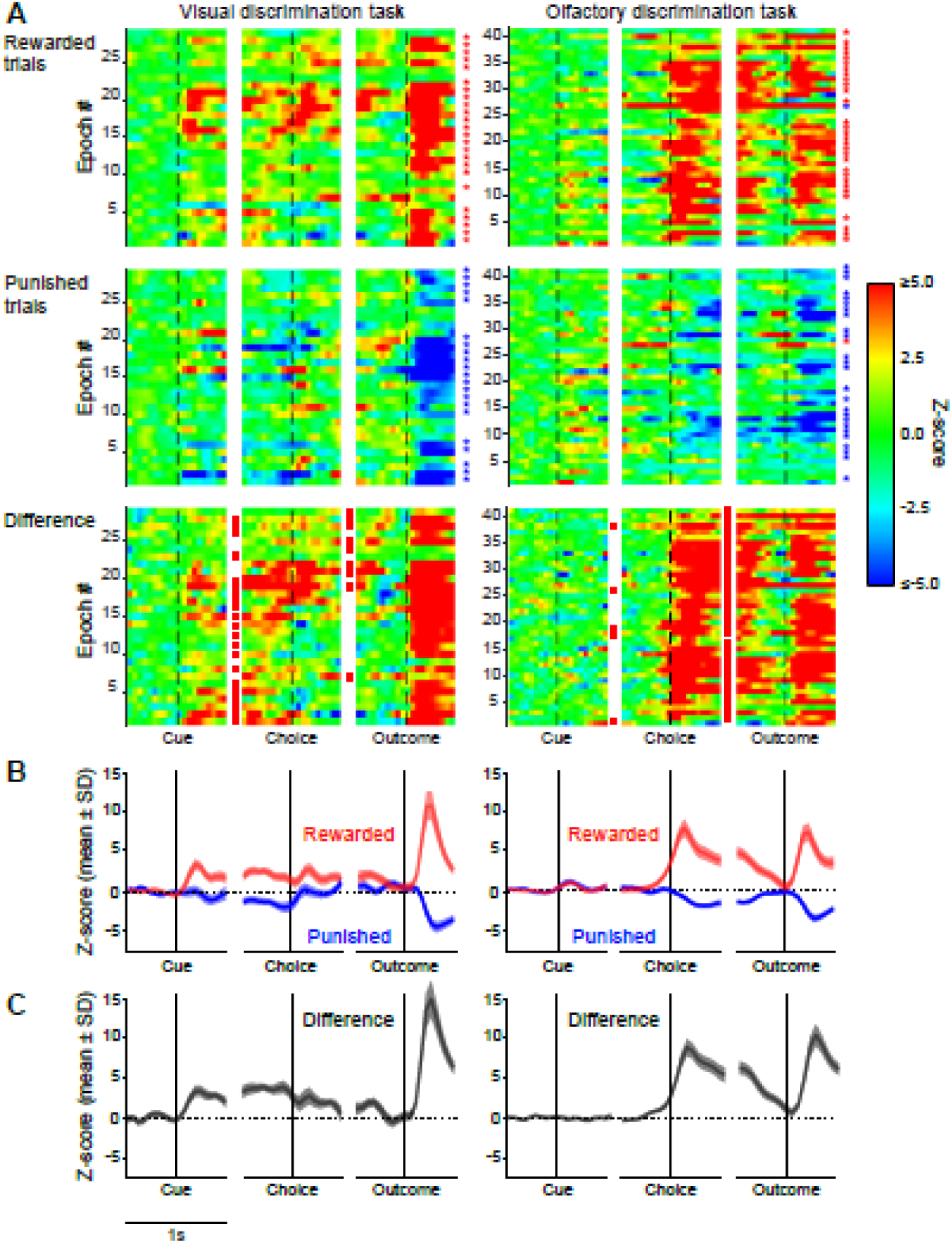
DA neurons with REW>PUN reward predicting firing rate increases. A, top two rows) The color rasters show z-scored firing rates in 0.5 s windows around the three principal task events (dashed vertical lines) for epochs with significant RP activity. “Cue” is when the mouse crossed the photobeam between the two compartments triggering the onset of visual and/or olfactory cues. “Choice” is the nose-poke. “Outcome” is the time of the onset of the outcome *signal* (feeder sounds for rewards or house light illumination for punishments). Each row represents firing of a neuron during a task epoch. The z-scores are calculated as [(firing rate of the pixel/mean firing rate of the baseline 0.5 s interval prior to cue onset of all trials in the epoch)/SD for the baseline period]. Red stars indicate significant increases in firing rate in the 0.5 s periods after reward or punishment (outcome) signals, relative to the 0.5 s baseline period prior to cue onset, while blue stars indicate significant firing rate decreases then (unpaired two-tailed t-test, p<0.05). A, third row) Reward prediction quantified as differences between values for rewarded and punished trials. Red squares indicate significantly higher firing rates in rewarded than in punished trials (unpaired two-tailed t-test, p<0.05) in the two respective task intervals. Staircase plots are ordered based on the latency of the peak firing rate between cue and outcome in the rewarded trials in the displayed data. The ±0.5 s windows may not cover the entire intervals between the task events, or they could overlap. Thus, there could be gaps or repetition of data in adjacent columns relative to the data from the actual interval used for the statistical analyses. B) Means (±SEM) of the data of the upper two rows of panels of A. C) Differences between the Rewarded and Punished traces in B.

**Figure 6.**
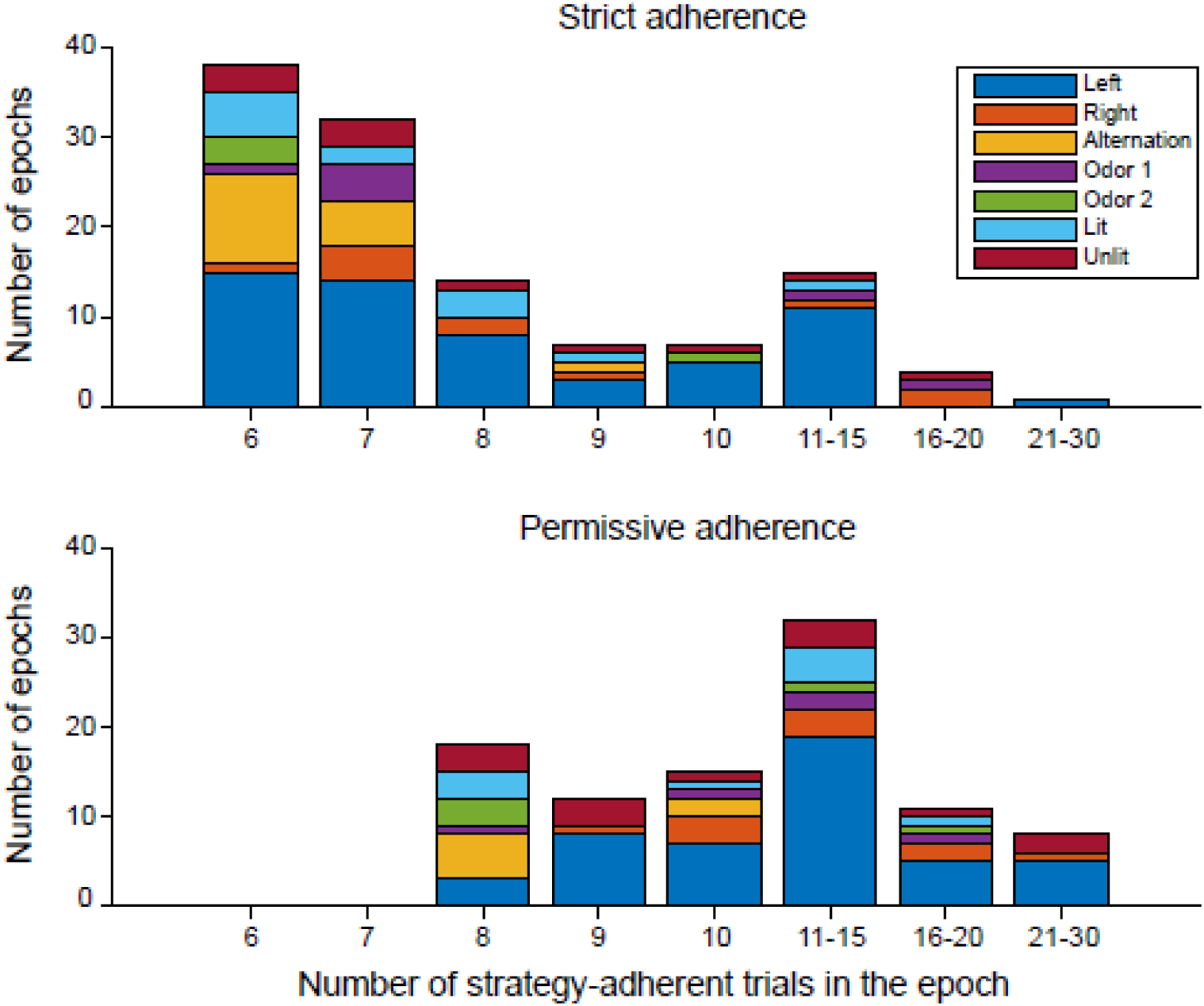
Incidences of series (“runs”) of successive trials adherent to strategies other than the currently rewarded rule in 53 epochs when DA REW>PUN or PUN>REW activity was recorded. (If more than one RP responding neuron was recorded within an epoch, the series was only counted once.) These tallies included 3201 trials. Strict and permissive criteria are as defined above. For the permissive adherence analysis (96 runs of trials), there are no runs of 6 or 7 trials since, by definition, adherent series had to be contained within sequences of at least eight trials.

DA neurons predicted whether the behavioral choice would be rewarded or punished according to the current rule even as the mice performed other strategies (Figure 5, 7 and 8). Of the 53 task epochs with DA reward-prediction activity (described below) and sufficient numbers of trials to analyze, 44 had series of trials that were performed according to strategies other than the current rule (according to the strict criterion of 100% adherence on successive trials of the series). While some of the strategies corresponded to rewarded task rules that the animals may have experienced (odor and light cues), others had never been rewarded even during pre-training (Left, Right and Alternation).

**Figure 7.**
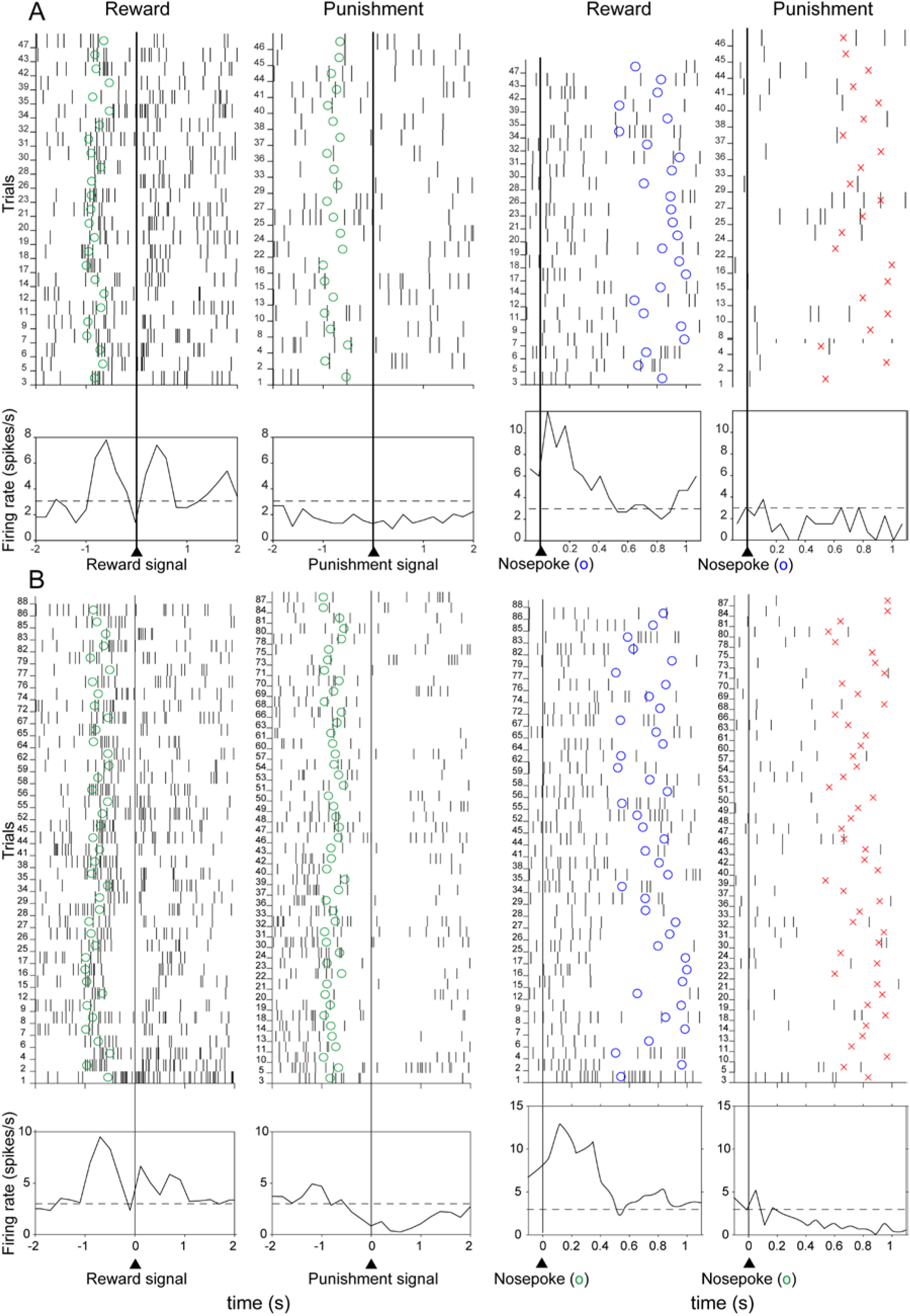
Examples of RP activity in DA neurons with one epoch. Green circles indicate the times of nose-poke choices in left two columns while at the right blue circles indicate timing of reward signals and red x’s indicate punishment signals. Horizontal dashed lines in the histograms below indicate background firing rate prior to trial onset. In the recording sessions in A and B, the task was complex (i.e., with distractor cues present). For A, the target cue was an odor, but the mouse chose the lit port with permissive adherence on trials 7-14, and 24-36. For the neuron in B, the target cue was an odor, and unrewarded strategies were: Alternation: trials 44-51; Lit port: trials 55-62; Left port: trials 75-88, all with permissive adherence. In both A and B, the RP response is significant in the [nose-poke, outcome signal] period, but the [cue onset, nose-poke] RP activity is only significant in A (p<0.05, unpaired t-test). The canonical activity RPE increase after reward signals is significant in both, but activity inhibition after the punishment signal is significant only in B (p<0.05, unpaired t-test). In this example, RP activity appear from the very first trial.

In no case did the activity clearly ramp up until trial outcome signals were presented. Rather, in Figure 5C, the Difference curves ramp downward. This contrasts with the upward ramping activity previously shown in dopaminergic neurons prior to rewards (e.g., Farrell et al, 2022), and which have been assimilated to a “motivational incentive” signal associated with reward seeking (Berridge and Robinson, 1998). Activity in the less numerous PUN>REW DA RP neurons are shown in Figure S4.

The proportions of RP activity in DA neurons occurring in epochs with simple vs. complex discriminations were not significantly different (17% vs 14%, χ^2^(1) = 0.73, p = 0.39; Table S1). There was a greater incidence of RP activity in epochs with olfactory target cues than visual target cues (20% vs 11%; χ^2^(1) = 8.4, p = 3.7 × 10^-3^; see Table S1).

Most of the REW>PUN RP activity in DA neurons were accompanied by canonical reward-prediction error (RPE) activity after the trial outcome signals. That is, reward signals triggered firing rate increases and punishment signals triggered reductions in firing rate (red and blue stars in Figure 5A, and Figures 7 and 8). Firing rates significantly increased after reward signals in 54/70 (77%) of DA neuron epochs (unpaired t-test, p<0.05) while firing rates decreased significantly after punishment signals in 47/70 (67%) (unpaired t-test, p<0.05). Note that when this firing rate reduction occurred, it only lasted at most for 1 s after punishment onset (Figures 7 and 8), even though the punishment period lasted 5 to 7 s. In Figures 7A and B, the similar numbers of rewarded and punished trials in each of these one-epoch sessions reflect the chance performance levels: 53% in (A) and 46% in (B). The trials with unrewarded strategies during the RP activity are indicated in the Legends of Figures 7 and 8.

**Figure 8.**
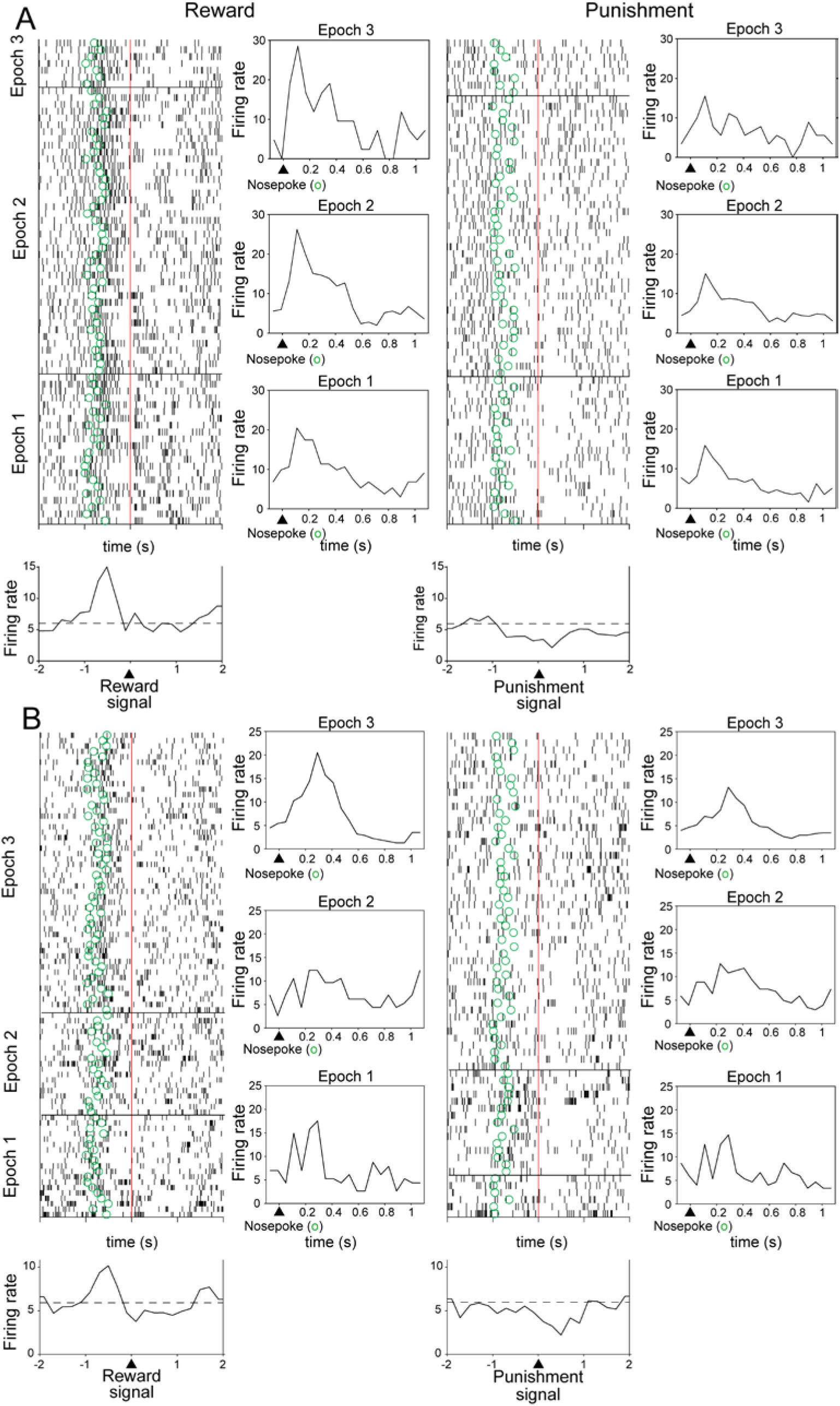
Examples of RP activity in DA neurons in sessions with multiple epochs. (Same format as Figure 7.) A) In epochs 1 and 3 the simple odor discrimination task was rewarded. In epoch 2, the complex odor task was rewarded. RP activity was significant in all three epochs (p<0.05, unpaired t-test). The mouse achieved criterion performance in epoch 1 only. Concerning RPE activity, inhibition was significant after punishment signals in epochs 1 and 2 only (p<0.05, unpaired t-test), but none had significant post-reward signal excitation (p>0.05, unpaired t-test). Performance levels were 46%, 51% and 47% in the respective epochs. Unrewarded strategies were: Alternation: trials 13-20 and 105-114; Left port: trials 21-29; Right port: trials 49-58 and 115-134; Unlit port: trials 59-67 and 84-95. B) In epoch 1 the task was the simple visual discrimination, in epoch 2, the complex visual task, and in epoch 3, the complex odor task. RP activity was significant only in epoch 3 (p<0.05, unpaired t-test). The mouse reached criterion performance in epochs 1 and 2. Concerning RPE activity, the inhibition after the punishment signal was significant in epoch 3 (p<0.05, unpaired t-test), but the firing rate was not significant higher after the reward signal (p>0.05, unpaired t-test). In epoch 3, the unrewarded strategy was Left nose-pokes on trials 122-130 and 137-150.

In the sessions of Figures 8A and B, the mouse reached criterion twice, and, accordingly, the task rule was changed each time. In Figure 8A, the RP response was significant in all three of the task epochs, while the performance level averaged only 51% correct. But in the neuron of Figure 8B, the RP response was significant only in the third epoch (while performance was 52%). For this reason and the fact that the animals were challenged by different tasks in the respective epochs, the analyses tally the incidence of RP activity for individual task epochs rather than over whole sessions (which could dilute significant effects present in only one epoch).

Note that RP activity could signal the reward contingency was visual or olfactory when the distractor cue was present (“complex” columns in Table S1, Figures 7 and 8). In ten sessions, DA RP neurons were recorded in successive epochs with visual and olfactory discriminations.

In five of these, RP activity was significant for only one of the sensory modalities, but in the other five, the neurons predicted reward in both tasks. Thus, the neurons did not necessarily code for correct choices exclusively for a single strategy policy. This would be consistent with the strategies being elaborated upstream from the DA neurons.

### Unidentified neuron activity

In the [nose-poke, outcome signal] interval, 10% of epochs for all tasks had REW>PUN activity (Table S1; Figure 8) and 7% were PUN>REW (Figure 9). However, in this interval for the odor discrimination task only, 16% had significant REW>PUN activity, and for the simple version of this task, this value rose to 23%. Note that in neurons in epochs with significant PUN>REW RP activity, there are many significant decreases in firing rate appearing after the onset of the olfactory target cue (Figure 9, cells 1-5, 7, 9 and 12) as well as after the onset of the visual cue. As for canonical RPE activity, firing rates significantly increased after reward signals in 40/58 (77%) of Unidentified neuron epochs while firing rates decreased significantly after punishment signals in 20/58 (34%; unpaired t-test, p<0.05). In a few cases, there was anti-canonical activity after outcome signals, with inhibition after reward signals and excitation after punishment signals (Figures 9 and 10).

**Figure 9.**
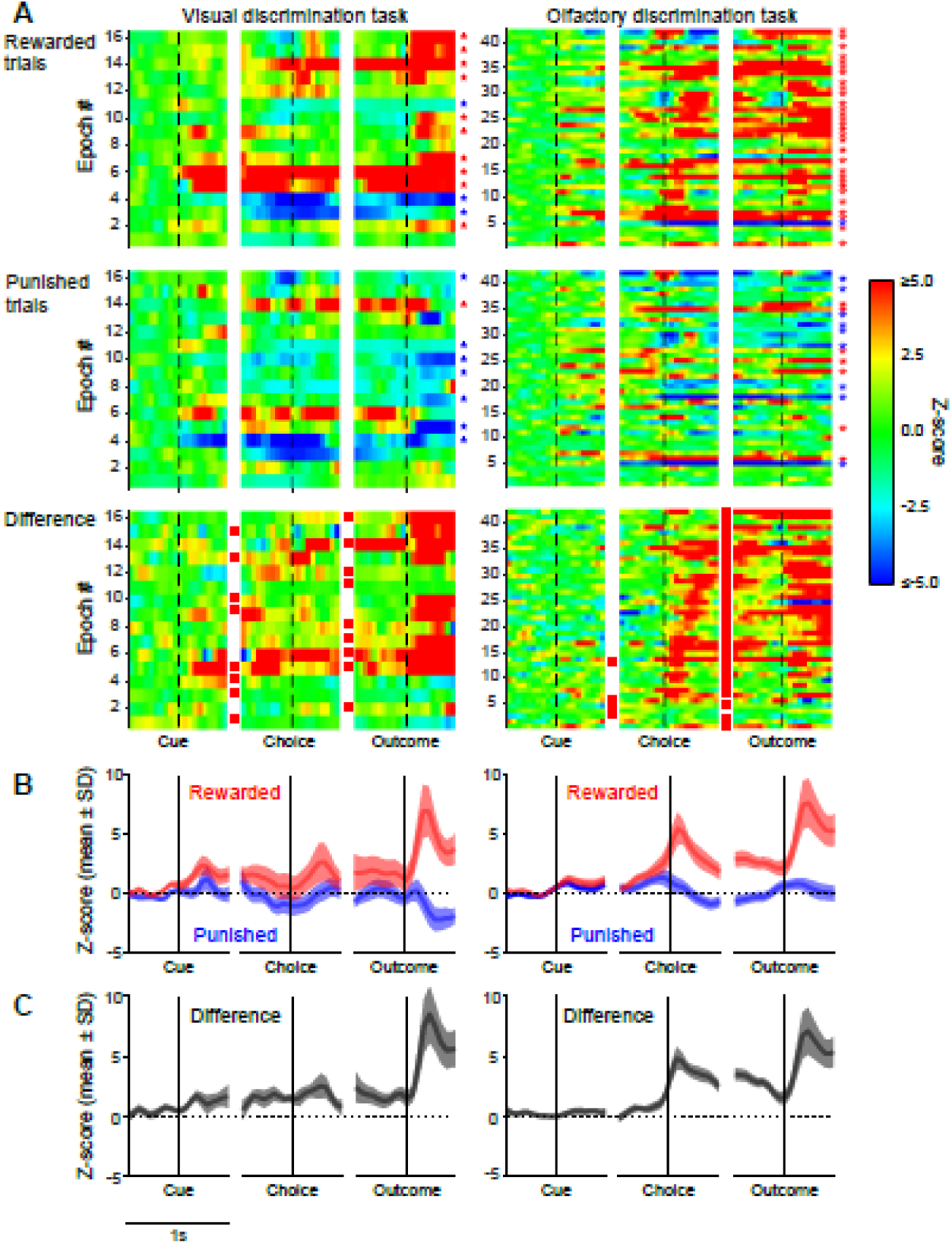
Unidentified neurons in epochs with REW>PUN RP firing rate increases. Same format as Figure 5.

**Figure 10.**
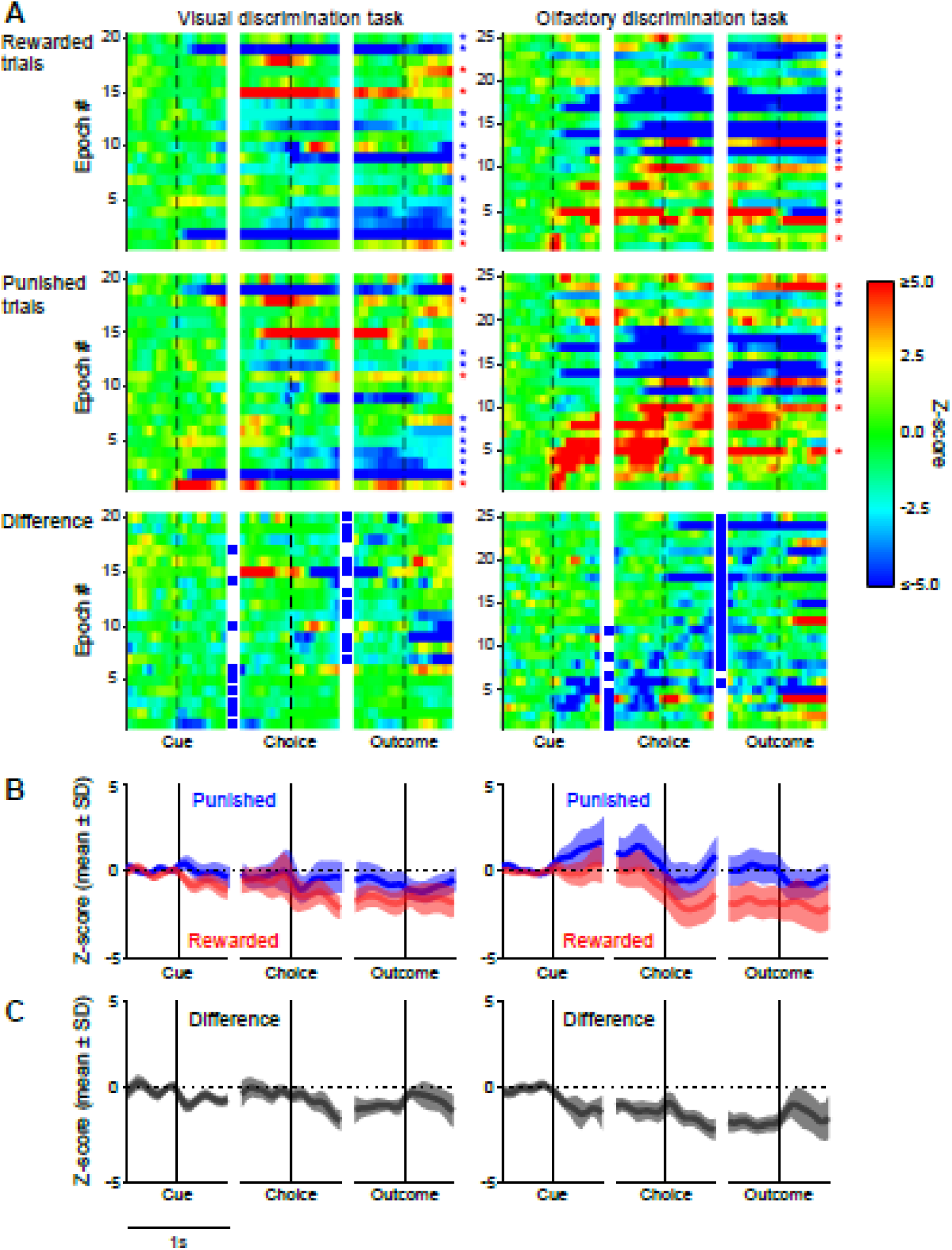
Unidentified reward predicting neurons with firing rate increases on punished trials (PUN>REW). Same format as Figure 5. Blue squares in the third row of A (“Difference”) mark neurons in epochs with significant PUN>REW activity in the respective intervals (unpaired two-tailed t-test, p<0.05).

### Behavioral correlates of pGABA neurons

In 87% of the pGABA neurons with RP epochs, the cells also increased firing rates prior to and following the nose-poke in the odor discrimination task. The firing rates in these epochs between cue onset and outcome signals were similar in rewarded and punished trials (Figure 11B, right; *r*= 0.984 p = 4.0 × 10^-11^, Pearson’s correlation of peak firing rate at Nosepoke) and thus the neurons were not reward predictive. As in the other cell categories, in the odor discrimination tasks the RP activity of pGABA neurons occurred in the [nose-poke, outcome] interval. PUN>REW RP activity in pGABA neurons are shown in Figure S5. The increased firing rates during nose-pokes in the pGABA neurons in both rewarded and punished trials corresponds to the activity profiles corresponding to the exertion of force previous found in GABAergic neurons (Lee et al., 2001; Jiang et al., 2025). It is not clear why the nose-poke activity increases did not occur in neurons with RP activity in the visual discrimination task, but this occurred in other pGABA neurons then. Figure 12 shows the activity of all pGABA neurons (including those with no RP activity) and some neurons have activity reductions during nose-pokes as well. Most of the pGABA neurons also have RPE activity (canonical and anti-canonical; cf. Stelzner et al., 2025)

**Figure 11.**
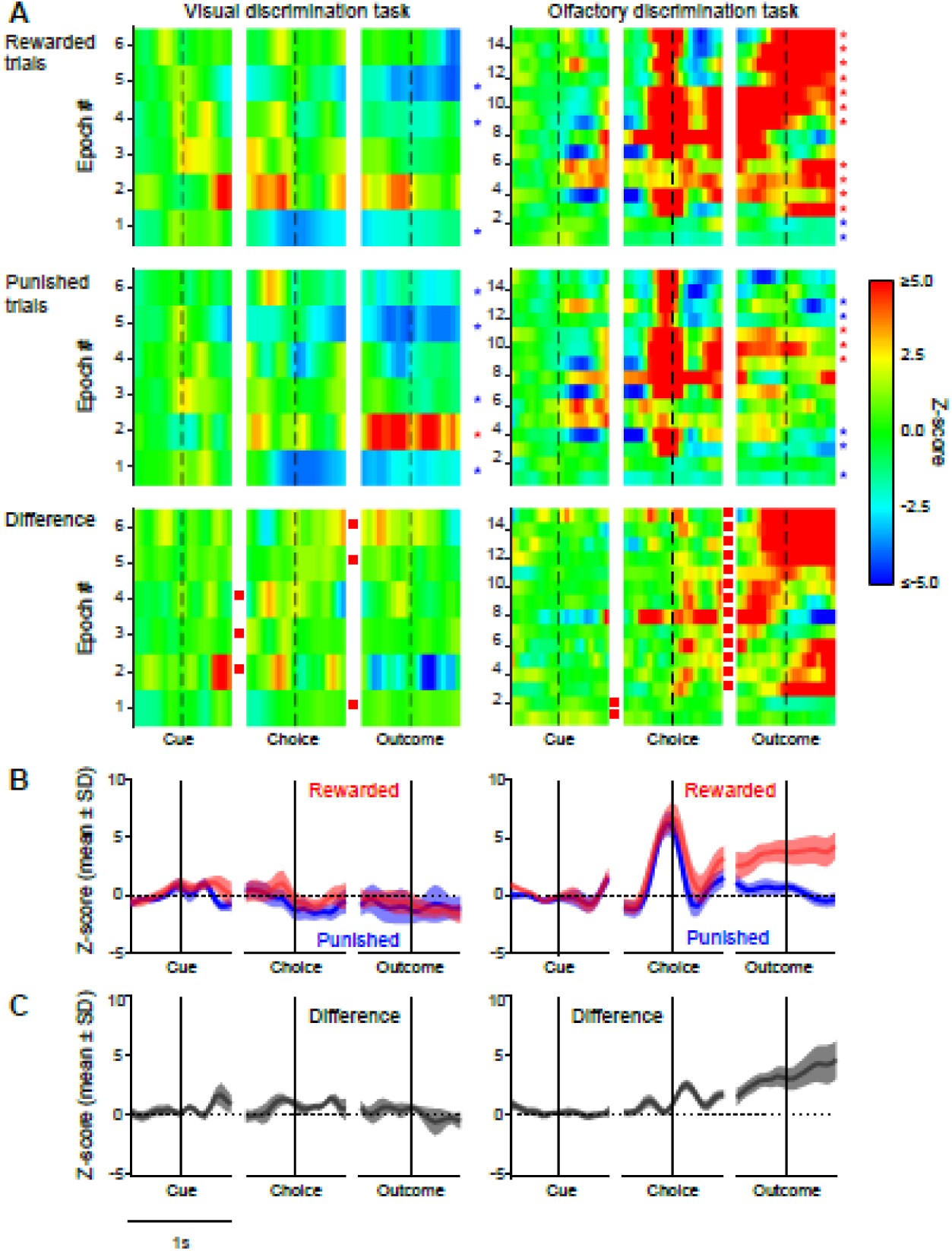
REW>PUN RP activity in pGABA neurons. Same format as Figure 5. A) In some epochs, significant intervals had relatively low magnitude z-scores (e.g., in “Difference”, in epoch 3 of the visual discrimination task and epochs 1 and 2 of the olfactory discrimination task) because of the high firing rates of these neurons.

**Figure 12.**
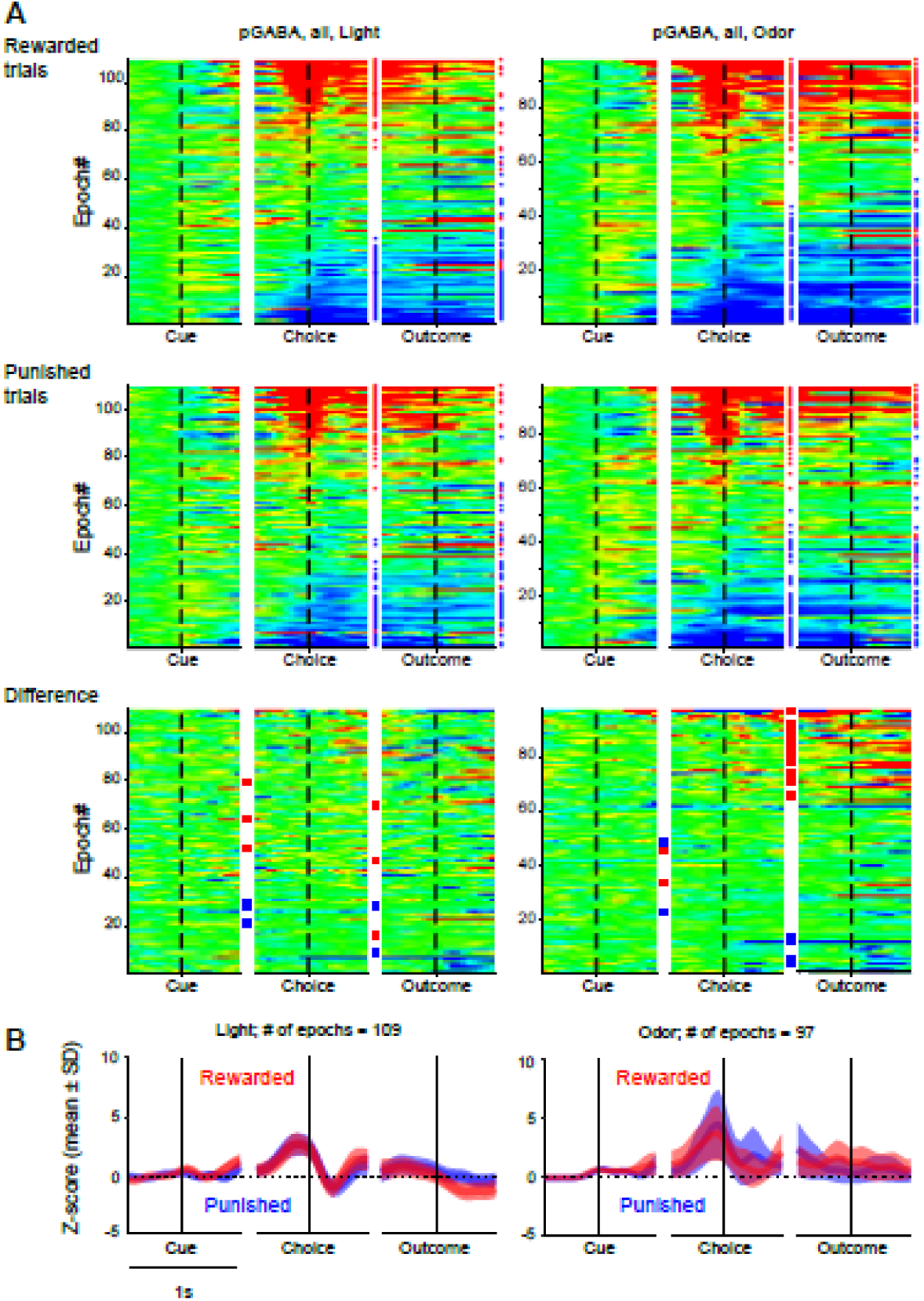
Activity of all pGABA neurons ordered by magnitude of nose-poke response magnitude. Same format as Figure 5 except that the columns of symbols between Choice and Outcome columns in A indicate nose-poke firing rate changes: significant increases (red stars) or decreases (blue stars) in firing rate during the interval [nose-poke – 0.5 s, nose-poke + 0.5 s] relative to baseline (unpaired two-tailed t-test, p<0.05). Stars to the right of the Outcome column indicate post-outcome firing increases and decreases as in previous Figures.

## Discussion

In summary, mice were challenged successively with self-paced visual and olfactory discrimination tasks with and without distractors and they executed unrewarded strategies prior to reaching criterion performance. During sub-criterion performance, some DA, pGABA and Unidentified VTA and SNc neurons fired phasically after decisions correctly predicting whether the behavioral choice would be rewarded or not, prior to emission of signals indicating the trial outcome. Thus, the neurons accurately predicted the imminent reward, even though the animal’s adherence to other strategies did not benefit from this information. The three cell categories had some differences in the incidence of response types. Furthermore, trial outcome signals (presented after a delay) evoked “reward prediction error” (RPE) activity that have been interpreted as the reward being a surprise (Schultz, 2016). A schematic model is proposed to explain these results in terms of a network for learning and decision-making with multiple modules sustaining alternative response policies.

### Sub-optimal strategies and their representations in the brain

In this task, as the mice adapted their behavior to changes in contingencies, behavioral choices were not random. Rather, the animals performed other strategies (often spatial) that were not currently, and often had never been, rewarded in this task. The Reinforcement Learning theory (Sutton & Barto, 2018) underlying some interpretations of midbrain dopaminergic neuron activity and later theoretical work emphasizing economic utility (Schultz, 2016) posit that behavior is optimized for increasing rewards at minimal cost and avoiding aversive stimuli. However, rats and mice have been observed to apply strategies that are sub-optimal in yielding reward relative to energy expended. Gruber and Hupta (2016) trained rats in a binary choice task with light cues where the optimal solution was programmed such that the rat received optimal rewards (50% of the time) if the animal’s choice of feeders was stochastic. Nonetheless, the rats performed a lose-shift strategy 68% of the time (but showed no significant tendency for win-stay strategy). Similarly, Molano-Mazón et al (2023) found that rats in a binary choice task selected suboptimal strategies after unrewarded trials. Srivathsa et al (2026) observed that aged rats alternate between optimal and sub-optimal strategies in a Morris water maze. Similar to our results, Ashwood et al (2022) analysed data from mice performing visual discrimination tasks and found that multiple non-optimal strategies were employed for blocks of tens to hundreds of trials. These strategies were manifested as hidden states in a generalized linear model, hidden Markov model (GLM-HMM) analysis of their data.

Cazettes, et al. (2023) observed mice as they performed multiple strategies in foraging tasks. They also recorded M2 neurons in these mice and found neural ensembles simultaneously encoding multiple strategies for these tasks. Thus, like the RP cells here, these neuron ensembles coded strategies not being currently employed by the animals. Note that M2 projects to VTA and SNc in mice (Watabe-Uchida, et al., 2012). Powell and Redish (2016) also found in rat prefrontal cortex neurons that transitions in strategy-related representations preceded behavioral strategy changes. This would be consistent with circuits containing different subsets of neurons, or “modules” underlying performance of respective strategies.

### Reward prediction activity prior to criterion performance in other brain areas

Schoenbaum et al (1998, 1999) found outcome predictive activity in basolateral amygdala (BLA) and orbitofrontal cortex neurons of rats during pre-criterion performance in a go/no-go olfactory discrimination task with a delay between onset of the behavioral choice and trial outcome signals. In most BLA neurons, firing rates increased for erroneous choices leading to punishment, opposite the polarity of most DA neurons here, but similar to others. After reversal of task contingencies, but prior to criterion performance, about half of the neurons no longer predicted trial outcome (Schoenbaum, et al., 1999), similar to our observations of RP activity not being maintained throughout all epochs of some recording sessions. The amygdalar activity also appeared at short latencies after choice onset. Midbrain dopaminergic neurons project to interneurons in basal amygdala (Brinley-Reed & McDonald, 1999; Pinard, et al., 2008; Lutas, et al., 2019; Tang et al., 2020). Furthermore, the VTA-amygdala-accumbens circuit acts as a positive feedback loop signaling positive experiences (Sun, et al., 2021). In mouse auditory cortex neurons, Drieu, et al (2025) also found a late component response in neuronal activity that predicted rewards in an auditory go/no-go discrimination task prior to when the animals achieved criterion performance.

### Novelty of this RP activity relative to dopaminergic activity in other operant tasks

While there are several studies of dopaminergic activity during instrumental learning and decision-making (e.g., Hollerman & Schultz, 1998; Bayer & Glimcher, 2005; Morris, et al., 2006; Nishino, et al., 1987; Phillips, et al., 2003; Roesch, et al., 2007; Roitman, et al., 2004; Stuber, et al., 2005; Jones et al., 2010), in many of these experiments, conditioned responses (CRs) and trial outcome signals were concurrent, and phasic firing rate increases after the CR could not be disambiguated from those related to the reward signal. Studies of dopamine release in the striatum and in recordings of midbrain dopaminergic neurons have often focused on animals undergoing classical conditioning or instrumental conditioning with instructed choices. Studies of instrumental conditioning with free choices have provided intriguing results pointing towards models other than actor/critic TD learning. Similar to the present results, Syed et al (2016) found that in an instrumental learning task, dopamine release in striatum was greater after initiation of correct than incorrect choice responses, but prior to the reward outcome signal. When the animals made the incorrect choice, the movement dynamics were slower than on rewarded trials, a possible confound. However, after correct choices, the DA release dynamics were fundamentally different from the present phasic activity: the dopamine levels gradually built up over the course of several seconds to peak after the trial outcome signal. These dynamics were also observed in recordings of VTA neurons under conditions of high uncertainty (Fiorillo, et al., 2003). These types of activity are generally assimilated with motivation incentive or drive for reward acquisition (Nishino, et al., 1987; Howe, et al., 2013), rather than with short-latency phasic firing rate increases related to motivation value for learning associations (Schultz et al., 1997; Wise, 2005). For example, in a task requiring self-initiated sequences of bar presses, dopamine release preceded and continued during these movements (Wassum et al., 2012). Again, the time course was over several seconds. There the firing rate increases were interpreted as representing incentive motivation.

The reward prediction activity here is phasic and on the same time scale as found in responses to rewards or to reward-associated cues Pavlovian and operant conditioning experiments, lasting for up to 200 ms (Schultz, 1997; Hollerman and Schultz, 1998). Lak et al (2016) distinguished a rapid time scale of phasic activation in DA neurons, at 0.1-0.2 s after cue presentation, but these corresponded to novelty-related activity which disappeared with familiarity. Subsequent activity, at 0.4-0.6 s after cue presentation, reflected learned reward value, but appeared only after performance levels had improved. Thus, this does not correspond to the activity found here. A third response longer time scale was characterized by sustained gradually ramping from cue presentation until reward delivery, similar to those discussed above. Again, this contrasts with the phasic RP activity here that immediately followed the behavioral response. For example, the RP activity here bears only superficial resemblance Goedhoop et al’s (2023) recordings of DA release in the nucleus accumbens (which receives VTA and SNc inputs) in animals performing similar Pavlovian and operant conditioning tasks. Ramping of DA release occurred only in their operant task, starting with the onset of the reward-predictive cue, and leading up to the operant response, similar to the observations of Lak et al (2020). This was interpreted as anticipation of a rewarded action. Ding et al (2025) found population activity of putative DA neurons of VTA was significant 15 to 18 trials after rule switches in their visual contrast discrimination task and 5 to 8 trials prior to the corresponding strategy changes. The RP activity of single units here was not restricted to these intervals.

Engelhard, et al (2019) recorded DA neurons in mice performing a visual discrimination task on a virtual T-maze. During the cue presentation, but prior to choice, some neurons selectively increased their discharge rate on trials when the mice made incorrect choices. Roesch et al (2007) found cue-triggered DA unit firing rate increases corresponded to the most valuable reward choice available, even on trials when the animal subsequently did not select this. After cues were turned off, but prior to movement for reward, the activity shifted to represent the actual choice. Thus, this is not similar to the observations here. Furthermore, cue selectivity developed with learning, while the activity here persisted over many trials, sometimes in epochs where learning did not occur prior to the end of the session.

Note that increased firing rates after aversive stimuli, an anti-canonical RPE activity, has been previously reported in midbrain DA neurons (Matsumoto and Hikosaka, 2009) as well as lateral habenula (Matsumoto and Hikosaka, 2007).

### Neuron identification

Since our primary aim was to better understand dopaminergic function, the dopaminergic neurons were optogenetically identified. It remains possible that the dopaminergic nature of some neurons in the Unidentified group was not detected for technical reasons. The putative GABAergic group was distinguished by waveform and firing rate criteria from the literature. However, there is no clear distinction between these three groups on the borders of these thresholds in Figure 3. In addition to possible false negatives for DA or GABA, it is possible that neurons in the Unidentified group could include glutamatergic neurons, those with GABA-glutamate co-transmission or possible sub-categories of neurons with co-transmitters. A greater proportion of Unidentified neurons had epochs with PUN>REW activity than the other two groups, and some of these could correspond to the type III VTA neurons of Cohen et al (2012) which decreased firing after presentation of odors predicting large rewards but not those predicting punishment. The latter neurons could not be identified as dopaminergic or GABAergic in that study. It would be interesting for future studies to determine whether certain response categories correspond to identified cell types such as glutamatergic, or specific co-transmitters with the others.

### Theoretical considerations

#### Dopaminergic activity and decision-making

The earliest RP activity appeared shortly after cue presentation for visual targets, but somewhat later when odor cues were targets, where they tended to occur after the nose-poke response. Since they reliably predicted reward or punishment the outcome of the behavioral choice must have already been determined then, and this could occur prior to the nose-poke. The earlier appearance of RP activity for the visual targets may be due to the visual cues being perceptible as soon as they were presented as the animals crossed into the chamber, while discriminating the odors may have improved with closer proximity to each of the nose-poke ports (and more time may have been required for the brain to process these signals).

This time frame is consistent with dopaminergic neuronal activity being associated with confidence in receiving a future reward as has been demonstrated by the canonical increase in DA neurons’ firing rates at the onset of cues that have been learned to predict reward (Hollerman and Schultz, 1998). Dopaminergic activity at the moment that a reward can be confidently predicted has also been found in a sensory discrimination task in a Y-maze. Benchenane et al (2010) observed that dopamine infusions in rat prefrontal cortex increased the coherence of hippocampal-prefrontal theta rhythms. This coherence also increased at the choice point of the Y maze on the trial when the rats successfully learned a new rule and thus could confidently anticipate receiving the reward at the end of the maze arm. The present work extends on this in showing that subsets of DA neurons can also anticipate whether the impending choice behavior will be rewarded, but here it is prior to when the learning is expressed in choice selection. Furthermore, this activity occurs at an appropriate time during training when they could ostensibly influence the network (including hippocampus and prefrontal cortex) toward selecting the rewarded response policy. A possible framework for underlying mechanisms for a role of REW>PUN RP activity in this is presented below.

#### Multiple strategies encoded in parallel “expert” networks

Acquiring optimal response policies in light of the history of rewards and punishments in a given situation can benefit from trial-and-error learning. In effect, the brain has to deduce the current response-outcome contingencies in a given situation, and select an action in the context of motivational priorities. These trial-and-error choices could be stochastic or based upon innate or previously learned strategies. The decisional arbitration process could then involve suppression of alternate action policies and/or promotion of beneficial ones. Doya (2002) reviewed early models composed of diverse experts and postulated a model for reinforcement learning with multiple modules, each containing a state prediction model and “responsibility signals”. There, arbitration is based upon a softmax estimate comparing prediction errors among the modules. Kurth-Nelson and Redish (2009) also proposed an arbitration process could involve multiple “micro-agents” enacted by brain circuits, each underlying different action policies. In Daw, et al. (2005), the brain engages a Bayesian mechanism to arbitrate between model-free and model-based processes. To resolve conflictual choices of these two systems, they use a reinforcement learning algorithm for learned optimal action control. This seeks to maximize accuracy by tracking the relative uncertainty of their respective predictions. Similarly, arbitration between experts (such as model-based vs model-free) is determined by the reliability of the respective systems in predicting reward (Lee et al, 2014; Shenhav, et al., 2013; O’Doherty et al., 2021). However, O’Doherty et al. (2021) stipulate that simpler experts (for example, for automatic habitual responses) may be preferred over ones requiring more deliberation and computation (see also Massi et al., 2022).

We propose a simple framework for the elaboration of REW>PUN RP activity and their role in learning and decision making in our task (see Figure 13), adapted from the Doya, et al (2002) and other models, some of which are cited above. Functional brain networks or “strategy modules” would correspond to respective behavioral policies. Such modules could include regions cited above as containing neurons coding alternative strategies: M2 cortex, amygdala, prefrontal cortex, orbitofrontal cortex, sensory cortices, as well as the basal ganglia loops (which include VTA/SNc) that they are connected with. In general, they could code innate strategies such as, in the present experiments, binary choices like go left, go right or spontaneous alternation. In other situations, this repertoire could extend to social interaction (including altruism), and security-related spatial preferences. Other innate strategies can be supramodal and could be informed by reward and punishment such as win-shift (e.g., foraging), win-stay, lose-shift (corresponding to depletion of resources). New strategies could be acquired by Pavlovian conditioning or goal-directed reinforcement learning from fortuitous trial-and-error choices, and then, later, can be added to the repertoire of strategies after sufficient repetition and reinforcement. There are two decisional arbitrators which receive decision propositions from a wide repertoire including stochastic, innate and acquired strategies. The *multipriority* arbitrator selects the behavioral choice, and is influenced by multiple factors including internal state (Onimus, et al, 2026), motivations, environmental context, level of impulsivity, trade-off between the satisfaction of performing habitual behaviors vs deliberating over a choice, etc (Figure 13). In contrast, the *optimizing* arbitrator has a single priority: selecting actions in order to optimize return versus energy cost while avoiding aversive outcomes.

**Figure 13.**
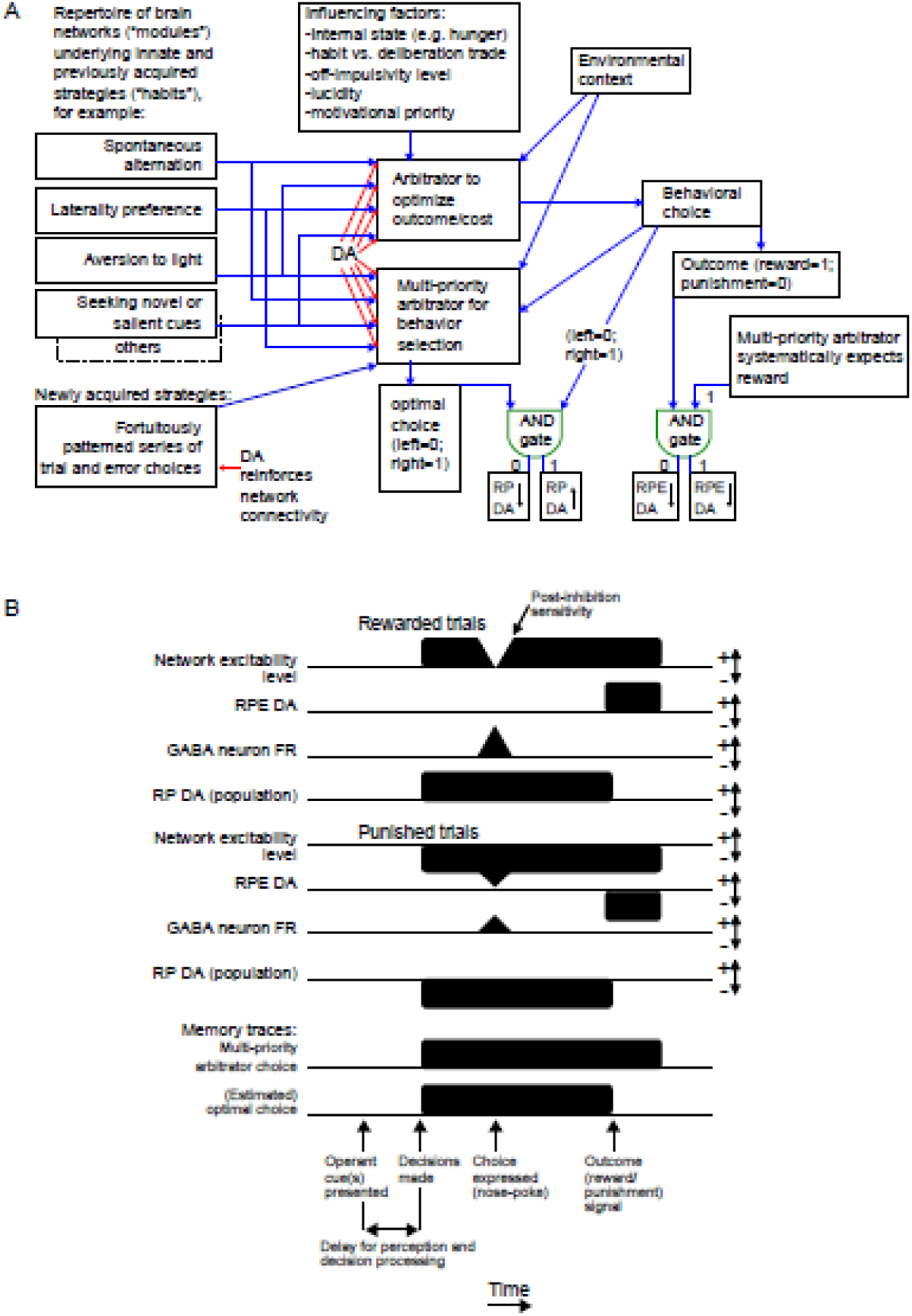
A framework engaging REW>PUN RP DA activity for the present task. A) The architecture of the framework. B) Timeline of the activity in the framework on a single trial. RP – reward prediction; RPE – canonical reward prediction error.

The optimizing arbitrator could operate with a TD reinforcement learning mechanism, taking into account the history of rewards, cues and actions in previous trials. Both arbitrators make their choice after an update of the environmental context. In the present tasks, this updating corresponds to perception and processing of the visual and/or olfactory cues (Figure 13B) and the decision can be made well before arrival at the decision point of the maze. Brain representations of upcoming (and previous) choices have been demonstrated in prospective (and retrospective) activity of hippocampal “splitter” cells (see e.g., Catanese, et al., 2014; the hippocampus is connected with VTA, e.g., Lisman & Grace, 2005). When behavioral choices coincide with the optimal choice, this would trigger the RP DA activity and dopamine release. But when they are different, this would trigger inhibition of DA neuron activity (AND gate to the left in Figure 13A). RPE processing is indicated by the AND gate to the right. DA neuron activation and inhibition would modify the strengths of the influence of the respective strategies on the behavioral choice arbitrator, and also reinforce or update the optimizing arbitrator to assure that it is indeed computing the optimal strategy. Note that in our binary choice task with distractors, only half of the trials would provide DA reward signal when a non-optimal strategy dominates the behavioral arbitrator. However, since rewarded trials would be concordant with the arbitrator deciding the optimal strategy, this would preferentially reinforce its connections on 100% of these trials (except when this strategy needs to be revised). In the case when the optimal choice arbitrator is no longer triggering regular rewards, a new strategy must be learned, the DA signal would reinforce the strengthening of synaptic connections that fortuitously led to high reward/punishment ratios, as in previous models. Over time, this new network would join the repertoire of strategy modules.

Note that the time course of downstream effects of dopamine are on the order of seconds (e.g. see Izhikevich, 2007; Gerstner et al., 2018; Gottschalk et al., 2025). As the nose-poke is made, the GABAergic neuron activity shown here would reduce spiking activity in the downstream network (Brown et al., 2012). (We assume that these the GABA neurons with nose-poke activity are projection neurons; Steffensen, et al., 1998.) This pause in activity could serve as a reset signal. In effect, network excitability and receptivity to RP DA signals would be heightened during post-GABAergic activity level rebounds. Furthermore, on trials when the optimal strategy is chosen, the corresponding module’s input to the behavioral choice arbitrator would have been recently active, and a spike-time dependent mechanism (Brzosko, et al., 2019) would permit the RP activity to selectively reinforce these connections (red arrows with “DA” in Figure 13A). Note that Lak et al (2020) found that optogenetic activation of VTA in DAT-Cre mice after stimulus presentation on 40% of trials selected randomly led to no change in behavior. This is consistent with the present framework which predicts that activation then would only be effective if selectively provided on trials corresponding to the optimal strategy, facilitating acquisition of new reward contingencies. RP activity would also prime connections for further reinforcement with the upcoming RPE reward-related DA activity. In this way, RP activity could provide a more timely and more effective critic signal than RPE signals alone. This DA signal would provide immediate feedback during decision-making and act gradually upon the network until the functional circuitry is modified sufficiently to lead to adopting a more adaptive behavioral choice policy.

## Acknowledgements

Thanks to France Maloumian for invaluable help with figures, to Yves Dupraz for constructing the experimental chamber, Estelle Anceaume for microscopy training, Dr. Michaël Zugaro for helpful suggestions and informatics support, Drs. Mehdi Khamassi, Benoît Girard, Céline Drieu and Linda Kokou for helpful comments, Dr. Guillaume Dugué and NeuroFabLab for facilitating 3D printing of headstages, Drs. Karim Benchenane and Liyang Xiang for help setting up the optical stimulation, and Dr. Philippe Faure and his team for advice on recording dopaminergic neurons with chronically implanted octrodes.

## Author contributions

JM and SIW designed and performed the experiments and obtained funding support. MNP performed experiments. JM designed the carrier for optic fibers and octrode driver assemblies, the optrodes and optogenetic stimulation apparatus, informatics control, and data acquisition systems and supervised constructing the experimental apparatus, with support from HN. MV and LV maintained and genotyped the mouse line and guided immunohistochemical processing. FP guided adaptation of the maze and behavioral protocols for training and recording. JM, VJO, and SIW analyzed the data with support from RT and HN, and wrote the manuscript. All authors approved of the manuscript.

## Ethics

All procedures were in accord with local (autorisation d’experimenter no. 75-1328-R; Comite d’Ethique pour l’Experimentation Animale no. 59, dossier 2012-0007) and international (European Directive 2010/63/EU; US National Institutes of Health guidelines) standards and legal regulations regarding the use and care of animals.

