## Supplementary Figures and Table for "A novel dopaminergic critic signal triggered by erroneous strategy choices in mice training in operant tasks": Supporting Figure legends.pdf

**Supporting Figure 1. Distribution of numbers of trials in epochs as a function of experience for all recording sessions.** Symbols are filled with red if the criterion performance to trigger changing the rule was achieved. Symbols occupying the nearby or same positions on the plot were displaced slightly for visibility. Symbol sizes reflect numbers of neurons with that property in a given epoch according to the numbers in the key.

**Supporting Figure 2. The distribution of reward prediction indices of all RP responsive neurons.** This index is calculated from the firing rates (FR) in the intervals as (rewarded FR - punished FR)/(rewarded FR + punished FR). Blue: REW>PUN; Red: PUN>REW, indicating that average firing rate during rewarded trials was significantly higher (or lower) than during the punished trials (unpaired t-test, p<0.05).

**Supporting Figure 3. Evolution of RP indices over the course of the epochs.** The RP indices are compared between the first and last six rewarded and punished trials of all epochs.

**Supporting Figure 4. PUN>REW RP activity in DA neurons.** Same format as Figure 7.

**Supporting Figure 5. PUN>REW RP activity in pGABA neurons.** Same format as Figure 7.

**Table S1. Relative incidence of RP responses in task epochs.** REW>PUN (or PUN>REW) signifies that the average firing rate was significantly higher (or lower) during rewarded trials than during the punished trials in this task interval (unpaired t-test, p<0.05). Epochs with REW>PUN RP responses in both intervals occurred in 6 DA neurons (3 for each target, 3 for complex and 3 for simple) and 2 Unidentified neurons (1 for each target, one simple, one complex). For NR>R RP responses in both intervals, there was one DA neuron (odor target, complex), one pGABA neuron (visual target, REW>PUN RP in the first interval and REW>PUN in the second, and two others had the inverse responses.
