## Supplementary Figures and Table for "A novel dopaminergic critic signal triggered by erroneous strategy choices in mice training in operant tasks": Supporting Table 1.pdf

### Cue-Choice interval

#### DA neurons

| | Visual target | | | Odor target | | | All targets | | |
|---|---|---|---|---|---|---|---|---|---|
| | Simple | Complex | Total | Simple | Complex | Total | Simple | Complex | Total |
| REW>PUN | 11 | 12 | 23 | 0 | 5 | 5 | 11 | 17 | 28 |
| PUN>REW | 1 | 1 | 2 | 0 | 4 | 4 | 1 | 5 | 6 |
| n.s. | 49 | 75 | 124 | 31 | 83 | 114 | 80 | 158 | 238 |
| Total | 61 | 88 | 149 | 31 | 92 | 123 | 92 | 180 | 272 |
| % REW>PUN | 18% | 14% | 15% | 0% | 5% | 4% | 12% | 9% | 10% |
| % PUN>REW | 2% | 1% | 1% | 0% | 4% | 3% | 1% | 3% | 2% |

#### pGABA neurons

| | Visual target | | | Odor target | | | All targets | | |
|---|---|---|---|---|---|---|---|---|---|
| | Simple | Complex | Total | Simple | Complex | Total | Simple | Complex | Total |
| REW>PUN | 2 | 1 | 3 | 0 | 2 | 2 | 2 | 3 | 5 |
| PUN>REW | 2 | 1 | 3 | 1 | 1 | 2 | 3 | 2 | 5 |
| n.s. | 38 | 65 | 103 | 20 | 73 | 93 | 58 | 138 | 196 |
| Total | 42 | 67 | 109 | 21 | 76 | 97 | 63 | 143 | 206 |
| % REW>PUN | 5% | 1% | 3% | 0% | 3% | 2% | 3% | 2% | 2% |
| % PUN>REW | 5% | 1% | 3% | 5% | 1% | 2% | 5% | 1% | 2% |

#### Unidentified neurons

| | Visual target | | | Odor target | | | All targets | | |
|---|---|---|---|---|---|---|---|---|---|
| | Simple | Complex | Total | Simple | Complex | Total | Simple | Complex | Total |
| REW>PUN | 6 | 3 | 9 | 4 | 1 | 5 | 10 | 4 | 14 |
| PUN>REW | 5 | 4 | 9 | 3 | 6 | 9 | 8 | 10 | 18 |
| n.s. | 82 | 145 | 227 | 54 | 176 | 230 | 136 | 321 | 457 |
| Total | 93 | 152 | 245 | 61 | 183 | 244 | 154 | 335 | 489 |
| % REW>PUN | 6% | 2% | 4% | 7% | 1% | 2% | 6% | 1% | 3% |
| % PUN>REW | 5% | 3% | 4% | 5% | 3% | 4% | 5% | 3% | 4% |

### Choice-Outcome interval

**DA neurons**

| | Visual target | | | Odor target | | | All targets | | |
|---|---|---|---|---|---|---|---|---|---|
| | Simple | Complex | Total | Simple | Complex | Total | Simple | Complex | Total |
| REW>PUN | 8 | 1 | 9 | 12 | 27 | 39 | 20 | 28 | 48 |
| PUN>REW | 0 | 2 | 2 | 2 | 3 | 5 | 2 | 5 | 7 |
| n.s. | 53 | 85 | 138 | 17 | 62 | 79 | 70 | 147 | 217 |
| Total | 61 | 88 | 149 | 31 | 92 | 123 | 92 | 180 | 272 |
| % REW>PUN | 13% | 1% | 6% | 39% | 29% | 32% | 22% | 16% | 18% |
| % PUN>REW | 0% | 2% | 1% | 6% | 3% | 4% | 2% | 3% | 3% |

**pGABA neurons**

| | Visual target | | | Odor target | | | All targets | | |
|---|---|---|---|---|---|---|---|---|---|
| | Simple | Complex | Total | Simple | Complex | Total | Simple | Complex | Total |
| REW>PUN | 2 | 1 | 3 | 1 | 12 | 13 | 3 | 13 | 16 |
| PUN>REW | 1 | 1 | 2 | 1 | 3 | 4 | 2 | 4 | 6 |
| n.s. | 39 | 65 | 104 | 19 | 61 | 80 | 58 | 126 | 184 |
| Total | 42 | 67 | 109 | 21 | 76 | 97 | 63 | 143 | 206 |
| % REW>PUN | 5% | 1% | 3% | 5% | 16% | 13% | 5% | 9% | 8% |
| % PUN>REW | 2% | 1% | 2% | 5% | 4% | 4% | 3% | 3% | 3% |

**Unidentified neurons**

| | Visual target | | | Odor target | | | All targets | | |
|---|---|---|---|---|---|---|---|---|---|
| | Simple | Complex | Total | Simple | Complex | Total | Simple | Complex | Total |
| REW>PUN | 5 | 4 | 9 | 14 | 26 | 40 | 19 | 30 | 49 |
| PUN>REW | 7 | 5 | 12 | 5 | 15 | 20 | 12 | 20 | 32 |
| n.s. | 81 | 143 | 224 | 42 | 142 | 184 | 123 | 285 | 408 |
| Total | 93 | 152 | 245 | 61 | 183 | 244 | 154 | 335 | 489 |
| % REW>PUN | 5% | 3% | 4% | 23% | 14% | 16% | 12% | 9% | 10% |
| % PUN>REW | 8% | 3% | 5% | 8% | 8% | 8% | 8% | 6% | 7% |
