## Supplementary figures and images for "A novel dopaminergic critic signal triggered by erroneous strategy choices in mice training in operant tasks"

### Figure S1 scatter plot.pdf

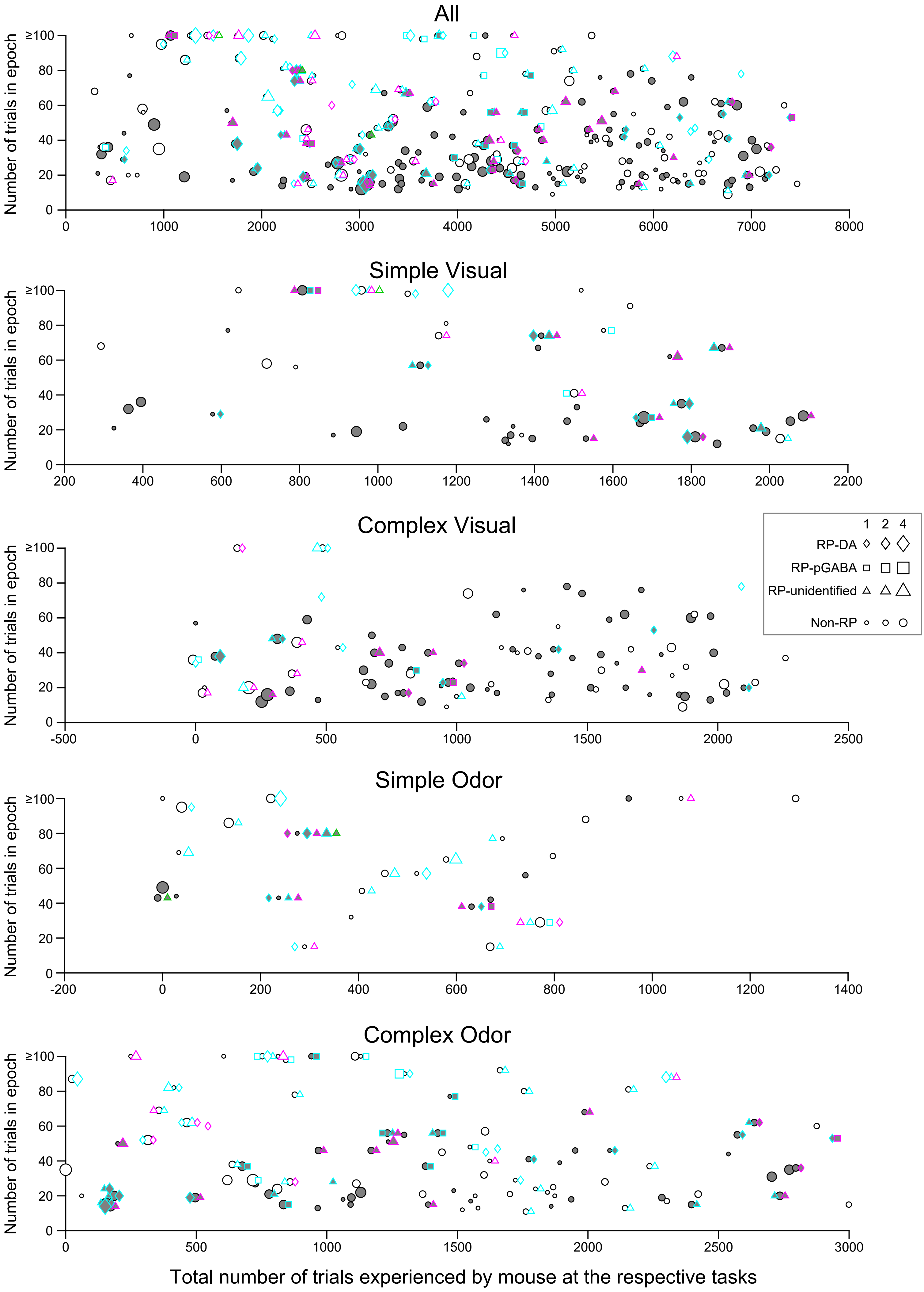


Cyan, REW>PUN; Magenta, PUN>REW; Green, Both; Gray shading, Criterion passed

### Figure S2 rp_index.pdf

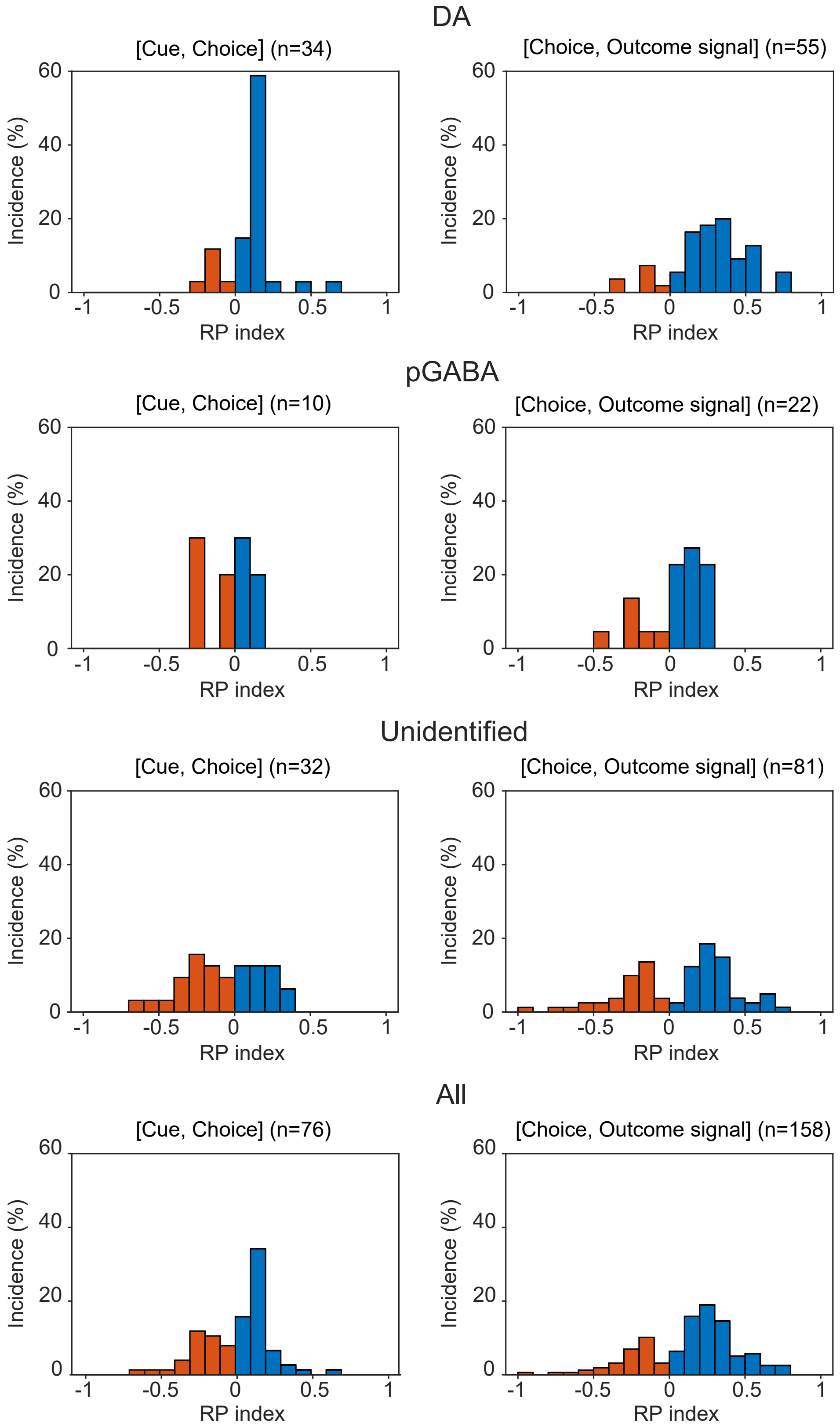

### Figure S3 rpindex_first_vs_last-Combined-v3.pdf

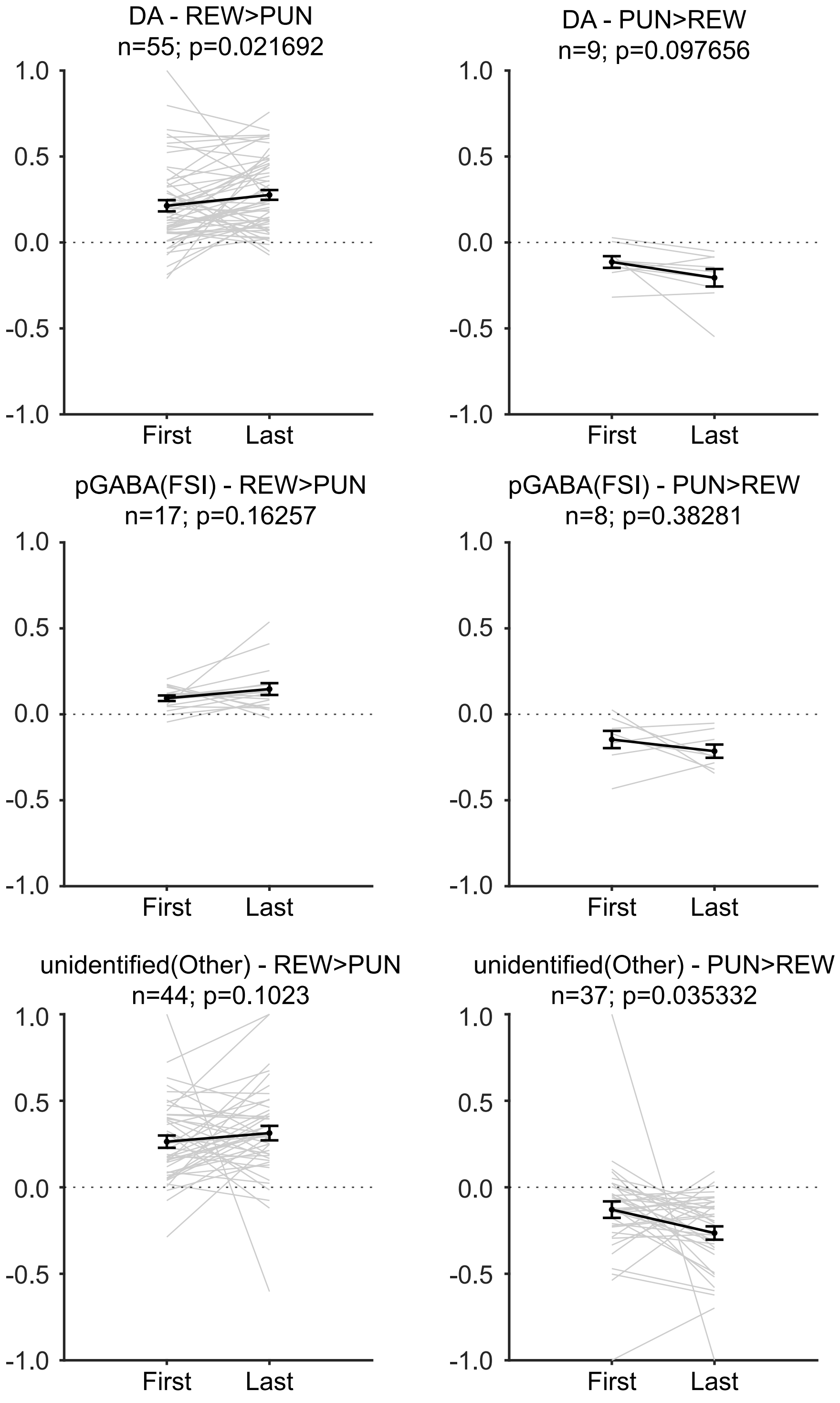

### Figure S4 da_rp_nr.pdf

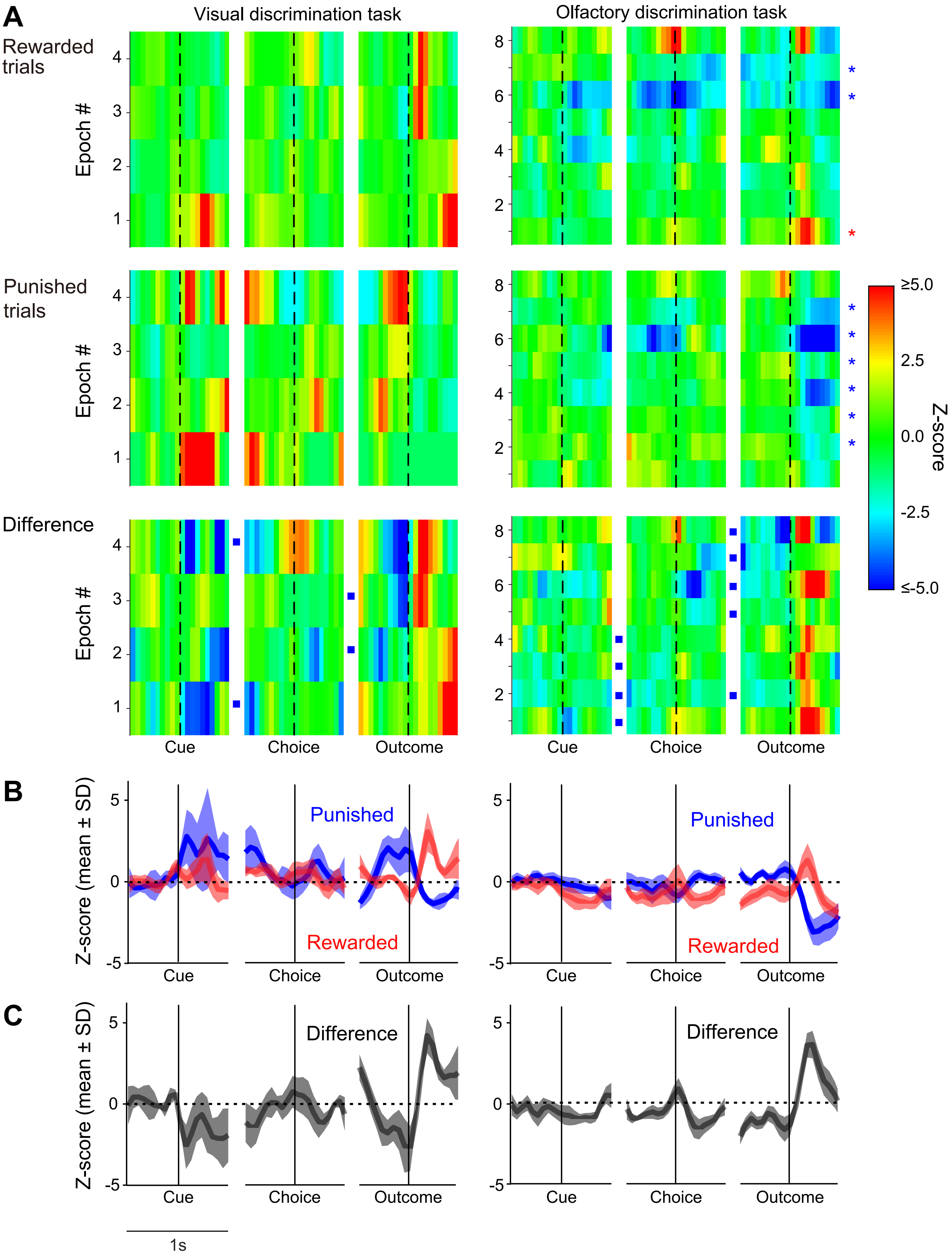

### Figure S5 2gaba_rp_nr-v6-import.pdf

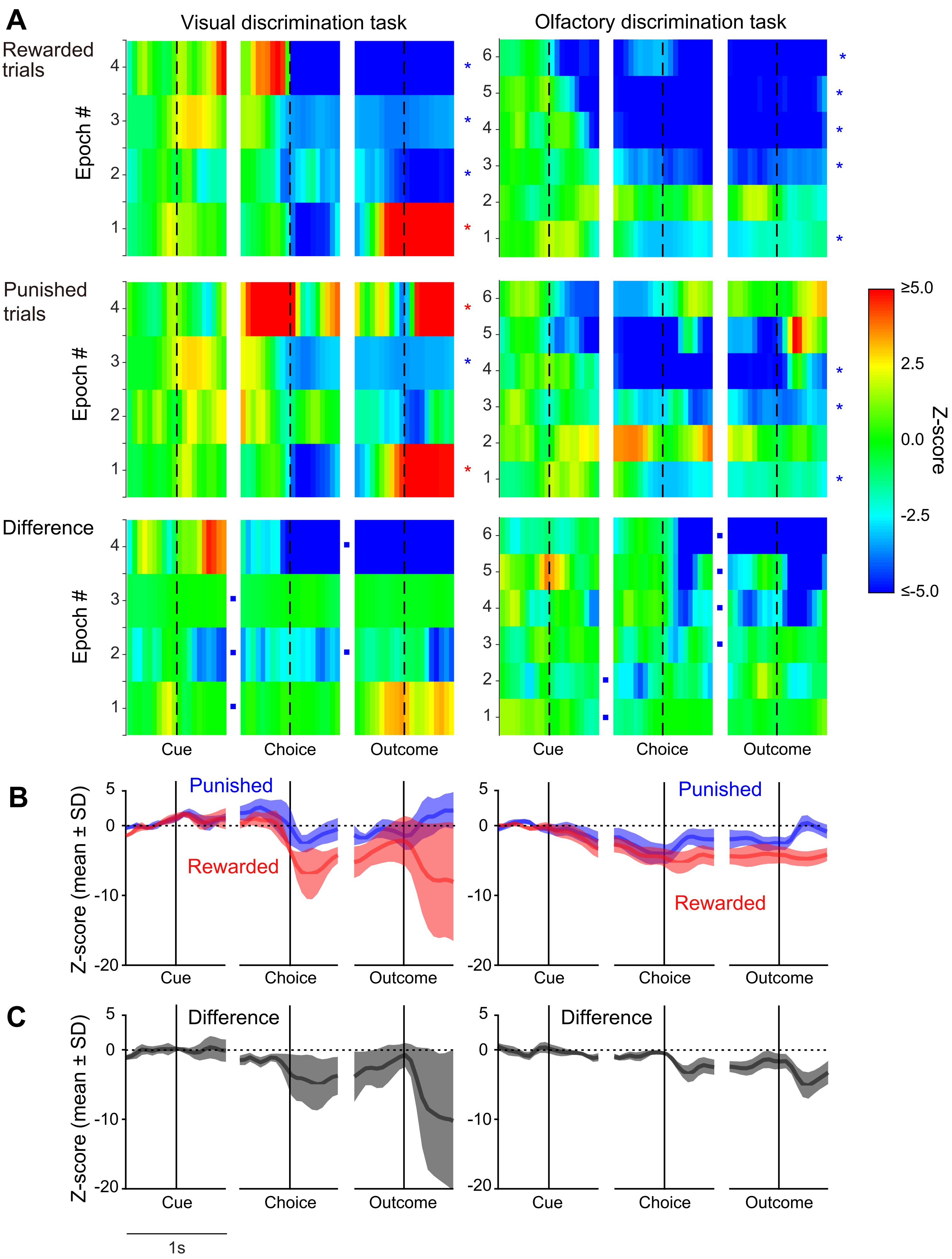
